# Stag1 and Stag2 regulate cell fate decisions in hematopoiesis through non-redundant topological control

**DOI:** 10.1101/581868

**Authors:** Aaron D. Viny, Robert L. Bowman, Yu Liu, Vincent-Philippe Lavallée, Shira E. Eisman, Wenbin Xiao, Benjamin H. Durham, Anastasia Navitski, Jane Park, Stephanie Braunstein, Elham Azizi, Matthew Witkin, Timour Baslan, Christopher J. Ott, Dana Pe’er, Job Dekker, Richard Koche, Ross L. Levine

## Abstract

Transcriptional regulators, including the cohesin complex member STAG2, are recurrently mutated in cancer. The role of STAG2 in gene regulation, hematopoiesis, and tumor suppression remains unresolved. We show Stag2 deletion in hematopoietic stem/progenitor cells (HSPC) results in altered hematopoietic function, increased self-renewal, and impaired differentiation. ChIP-sequencing revealed that while Stag2 and Stag1 can bind the same loci, a component of Stag2 binding sites are unoccupied by Stag1 even in Stag2-deficient HSPCs. While concurrent loss of Stag2 and Stag1 abrogated hematopoiesis, Stag2 loss alone decreased chromatin accessibility and transcription of lineage-specification genes, including Ebf1 and Pax5, leading to blunted HSPC commitment to the B-cell lineage. Our data illustrate a role for Stag2 in transformation and transcriptional dysregulation distinct from its shared role with Stag1 in chromosomal segregation.

**One Sentence Summary:** Stag1 rescues topologically associated domains in the absence of Stag2, but cannot restore chromatin architecture required for hematopoietic lineage commitment

## Main text

Cell-type specific transcriptional programs are regulated in part by the activity of tissue-specific transcription factors and enhancer elements within structurally defined topologically associating domains (TADs) (*1-3*). The genes which contribute to transcriptional regulation, including members of the cohesin complex, are frequently mutated in human cancers, including leukemias(*4-9*). Cohesin is a tripartite ring comprised of three structural proteins SMC1A, SMC3, and RAD21; this core complex binds to either STAG1 or STAG2. *STAG2*, which is an X-linked gene, is the most commonly mutated cohesin complex member in human cancer, with recurrent nonsense, frameshift, and missense mutations in Ewing’s Sarcoma (40-60%)(*9*), bladder cancer (20-35%)(*7, 8*), glioblastoma (4-5%)(*7*), myelodysplastic syndrome (MDS)(5-20%)(*5, 6*), and acute myeloid leukemia (AML)(2-12%)(*5*). This is in contrast to SMC1A, SMC3, and RAD21, which are more commonly mutated in myeloid malignancies than in a broad spectrum of human cancers. The cohesin complex is essential in pleiotropic cellular and gene regulatory functions, including chromosome segregation(*10, 11*) and in the formation of DNA loops(*3, 12-14*) which regulate gene expression(*13, 15, 16*). However, the high mutational frequency and unique mutational spectrum of *STAG2* in human cancers(*5-7*) suggests a distinct role in homeostasis and tumor suppression(*17, 18*) which has not been delineated.

## Results

### Loss of Stag2 alters hematopoietic stem cell function

In order to elucidate the role of *Stag1* and *Stag2* in hematopoietic function/transformation, we generated *Stag1* and *Stag2* conditional knockout alleles. We induced somatic deletion of *Stag1* or *Stag2* in hematopoietic cells through *Mx1*-Cre mediated excision (Figure S1A-H). Following polyinosinic:polycytadinic (PIPC)-mediated deletion, *Mx1*-cre *Stag2* ^−/−^ mice compared to cre-negative *Stag2*^fl/fl^ (referred henceforth as KO and WT respectively) manifested an increase in hematopoietic stem cells (HSC), including a significant (2.2-fold; p<0.01) increase in the proportion of LSK (Lineage^−^Sca1^+^cKit^+^) cells (Figure 1A). *Stag2* loss, but not Stag1 loss, induced an expansion of multipotent progenitors (MPP: LSK^+^Cd150^−^Cd48^+^Cd127^−^; p<0.04), short term HSCs (ST-HSC: LSK^+^Cd150^−^Cd48^−^; p<0.001), and long-term HSCs (LT-HSC: LSK^+^Cd150^+^Cd48^−^; p<0.02) (Figure S2A). We also observed a significant increase in granulocyte macrophage progenitors (GMP: Lineage^−^cKit^+^Sca1^−^Cd34^+^Fcγ^+^; p<0.001; Figure S2B-C) in the *Stag2-*deficient hematopoietic compartment.

**Fig 1.**
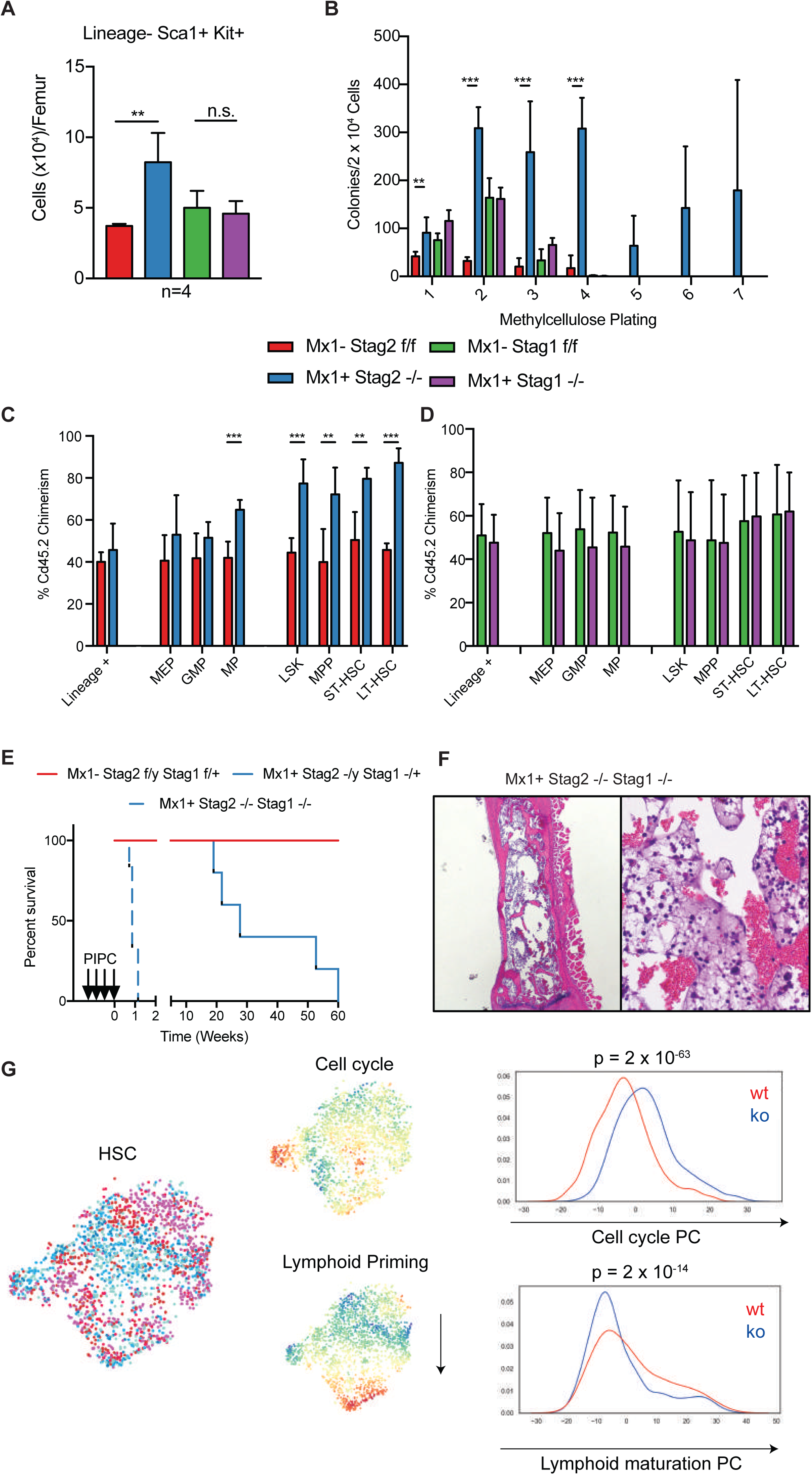
Hematopoietic specific loss of Stag2, but not Stag1, results in altered stem cell function. **A)** Stag2 KO, but not Stag1 KO mice have increased LSK (Lin^−^Kit^+^Sca1^+^) hematopoietic stem cells (Log_2_FC=2.2; p<0.01) **B)** Whole bone marrow plated in cytokine-enriched methylcellulose reveals that Stag2 KO marrow has increased self-renewal capacity and can serially replate, but WT and Stag1 KO cannot. Competitive bone marrow transplantation of **C)** Stag2 WT (Red)/KO (Blue) or **D)** Stag1 WT (Green)/KO (Purple) bone marrow mixed 1:1 with Cd45.1 normal marrow shows increased chimerism in stem and progenitor populations in the bone marrow at 16 weeks. **E**) Kaplan-Meier curve shows Stag2/Stag1 is lethal with a median survival of 0.7 weeks (p=0.01), while Stag2 KO/Stag1^−/+^ has a lethal phenotype with a median survival of 27.7 weeks (p=0.05). **F**) Hematoxylin and eosin (H&E) staining of Stag2/Stag1 KO bone marrow reveals marked cellular aplasia. **G**) Left: t-SNE projection of library-size normalized and log transformed data for inferred HSC subset (2025 cells). Each dot represents a single cell colored by genetic condition (Stag2 KO: shades of blue, Stag2 WT: shades of red). Middle: t-SNE projection colored by the second and third principal components most correlated with lymphoid priming and cell cycle, respectively. Right: Distribution of cells along lymphoid and cell cycle components; p-values for Mann-Whitney U test are shown.

We next investigated the role of Stag2 and Stag1 in hematopoietic self-renewal. *Stag2*-deficient hematopoietic cells displayed increased serial replating capacity in methylcellulose culture assays (≥7 platings), whereas WT and *Stag1* KO cells were not able to form colonies beyond 3 platings (Figure 1B). We next performed competitive transplantation assays to assess the impact of Stag2/Stag1 loss on self-renewal *in vivo*. We observed reduced *Stag2-*deficient derived chimerism in the peripheral blood of mice reconstituted with Stag2 KO and WT competitor cells (p<0.001; Figure S2D-F). However, within the bone marrow of transplant recipient mice, we found a significant increase in *Stag2* KO derived chimerism, which was most significant in the LSK compartment (p<0.001) including in LT-HSCs (p<0.001; Figure 1C) but not in lineage positive cells (p≤0.37). *Stag1* loss did not impact *in vivo* self-renewal or differentiation output (Figure 1D; S2G-I). These data suggest that *Stag2* loss induces both an increase in self-renewal and reduced differentiation capacity in HSCs, which are critical features of hematopoietic transformation.

To determine if Stag2 loss was associated with the expansion of hematopoietic cells with clonal cytogenetic alterations, we performed karyotype analysis in *Stag2* and *Stag1* WT and KO bone marrow cells. Chromosomal aberrations by metaphase karyotyping were not detected in either *Stag2-* or *Stag1-*deficient hematopoietic cells (n=7; Figure S3A-B). We did observe an increase in the proportion of cells with premature sister chromatid separation in *Stag2* KO bone marrow (Figure S3C); however, consistent with metaphase cytogenetics, we did not observe clonal copy number alterations on low-pass whole genome sequencing (Figure S3D-E). This data suggested that the increase in hematopoietic self-renewal induced by *Stag2* loss is not due to genomic instability and resultant clonal evolution.

### Stag2 and Stag1 co-deletion is synthetic lethal

The observation that *Stag2-*deficient HSCs displayed increased self-renewal is in contrast to the lethality observed with loss of the core cohesin component *Smc3* in hematopoietic cells(*19*). We next sought to evaluate potential compensatory function among the Stag paralogs. We found *Stag1* transcript and protein expression was significantly increased in *Stag2* KO hematopoietic cells, whereas *Stag2* expression was not altered in *Stag1* KO hematopoietic cells (Figure S4A-D). Analysis of the AML TCGA dataset showed that AML patients with STAG2 mutations expressed higher levels of *STAG1* (p<0.006; Figure S4E), suggesting compensatory regulation. We hypothesized that Stag1 and Stag2 may have redundant roles with respect to chromatid segregation such that Stag2 loss does not fully abrogate cohesin function due to residual Stag1 function. *In vitro* studies have recently nominated STAG1 as a synthetic lethal target in *STAG2* mutant cell lines(*20*); however, this has not been assessed *in vivo*. We generated mice with hematopoietic cells retaining one or zero copies of *Stag1* on the background of a *Stag2* (hemizygous male) KO genotype. *Stag2*^*-/y*^*/Stag1*^*+/-*^ mice developed rapid, progressive thrombocytopenia (p<0.001) and impaired survival (p<0.001) due to bone marrow failure (median survival of 27.7 weeks). Complete loss of *Stag1* and *Stag2* in hematopoietic cells induced rapid lethality (median survival 0.7 weeks; Figure 1E; Figure S4F). *Stag1/Stag2* deficient mice were characterized by pancytopenia and bone marrow aplasia, consistent with redundancy in hematopoietic function and in alignment with the absolute requirement for core cohesin complex members (Figure 1F). Similar to mice with bi-allelic deletion of *Smc3*, we observed marked chromosomal alterations on metaphase spreads of the *Stag1/Stag2* deleted bone marrow (Figure S4G). Collectively, these data suggest a role for Stag2 in gene regulation and tumor suppression that is not shared with Stag1, whereas they have a shared, redundant role in chromosomal segregation.

### Stag2 alters transcriptional lineage commitment

We hypothesized that *Stag2* loss induced alterations in transcriptional output which dysregulate hematopoietic function and promote transformation. To investigate transcriptional mechanisms regulating self-renewal and differentiation in the context of *Stag2* loss, we performed RNA-sequencing on *Stag2* KO and WT LSK cells. We identified 186 genes which were significantly downregulated and 42 genes which were significantly upregulated in *Stag2* KO compared to WT cells (Figure S5A). We observed reduced expression of *Ebf1, Vbrep3, Cd19, Bank1*, and *Pax5*, suggesting a loss of B-cell lineage commitment(*21, 22*) in *Stag2-*deficient stem/progenitor cells. We observed reduced expression of *Vwf(23), Hba-a1, Hba-a2, Hbb-b1(24), Fcgr2b(25), Ccr2(26), Spib(27), S100a8, and S100a9(28)*, genes associated with myeloid/erythroid lineage commitment. Gene set enrichment analysis (GSEA) revealed that Stag2 deficient LSKs showed negative enrichment of genes associated with cell lineage specification (Figure S5B). We further found increased expression of the Hoxb4-target *Hemgn(29)*, as well as of *Itgb3*(*30*), a Periostin-binding integrin shown to mediate HSC self-renewal. Coupled with downregulation of *Flt3(31)* and *Selp*(*32*) (P-selectin), these findings indicate an LSK pool that is transcriptionally skewed towards increased self-renewal.

Given the heterogeneity of the LSK compartment, we sought to determine if the transcriptional alterations observed in bulk LSKs were due to a population wide shift in expression or a result of differential expression within a subset of LSKs. We performed single cell RNA-sequencing (scRNAseq) of ∼24,000 Lin^−^ hematopoietic stem/progenitor cells (HSPC) from Stag2 KO (n=3) and WT (n=3) mice (cohort annotations in Figure S5C-E). We found that within the HSC population, representing ∼8% of cells, that the compartment distribution was skewed with KO HSCs having enriched expression of cell cycle gene signatures (p<10^−63^; Figure 1G, S5F top panel). Concomitantly, these HSC showed decreased expression of early lymphoid commitment signatures (p<10^−14^; Figure 1G, S5F bottom panel) even at the earliest HSC states, consistent with negative enrichment for cell specification. We expanded upon these findings by comparing bulk LSK differentially expressed genes to gene expression data from dormant and active HSCs separately isolated through genetic lineage tracing studies(*33*). We observed that genes that were decreased in expression in *Stag2*-deficient mice were most highly expressed in slow-cycling quiescent HSCs, while the upregulated genes were most highly expressed in the proliferative MPP1 fraction (LSK, Cd34^+^, Flt3^−^) (Figure S5G). Similar findings were observed when comparing our data to single cell RNA-seq data, such that the most downregulated genes in *Stag2-*deficient LSKs were most highly expressed in the most quiescent HSCs (Figure S5H). Collectively, these analyses highlight a transcriptional signature associated with decreased lineage commitment, decreased quiescence, and enhanced self-renewal in the setting of STAG2 loss.

### Stag2 loss impairs differentiation and induces dysplasia

To determine the consequences of these transcriptional alterations *in vivo*, we assessed the phenotype of mice with *Stag2*-deficient hematopoiesis over time. Hematopoietic-specific *Stag2* loss resulted in progressive leukopenia (p<0.02) and thrombocytopenia (p<0.01; Figure S6A). We observed an expansion in myeloid cells and a significant reduction in B lymphocytes (p<0.001; Figure 2A) in the peripheral blood, which was not seen with *Stag1* deletion. *Stag2* loss, but not *Stag1* loss, induced morphologic alterations including an increase in immature myeloid cells, an expansion of small hypolobated megakaryocytes, and nuclear:cytoplasmic dyssynchrony in the erythroid lineage (Figure 2B). These abnormalities are features of human myelodysplasia—consistent with the observation that 5-15% of MDS patients have loss-of-function mutations in *STAG2*(*5, 6*).

**Fig 2.**
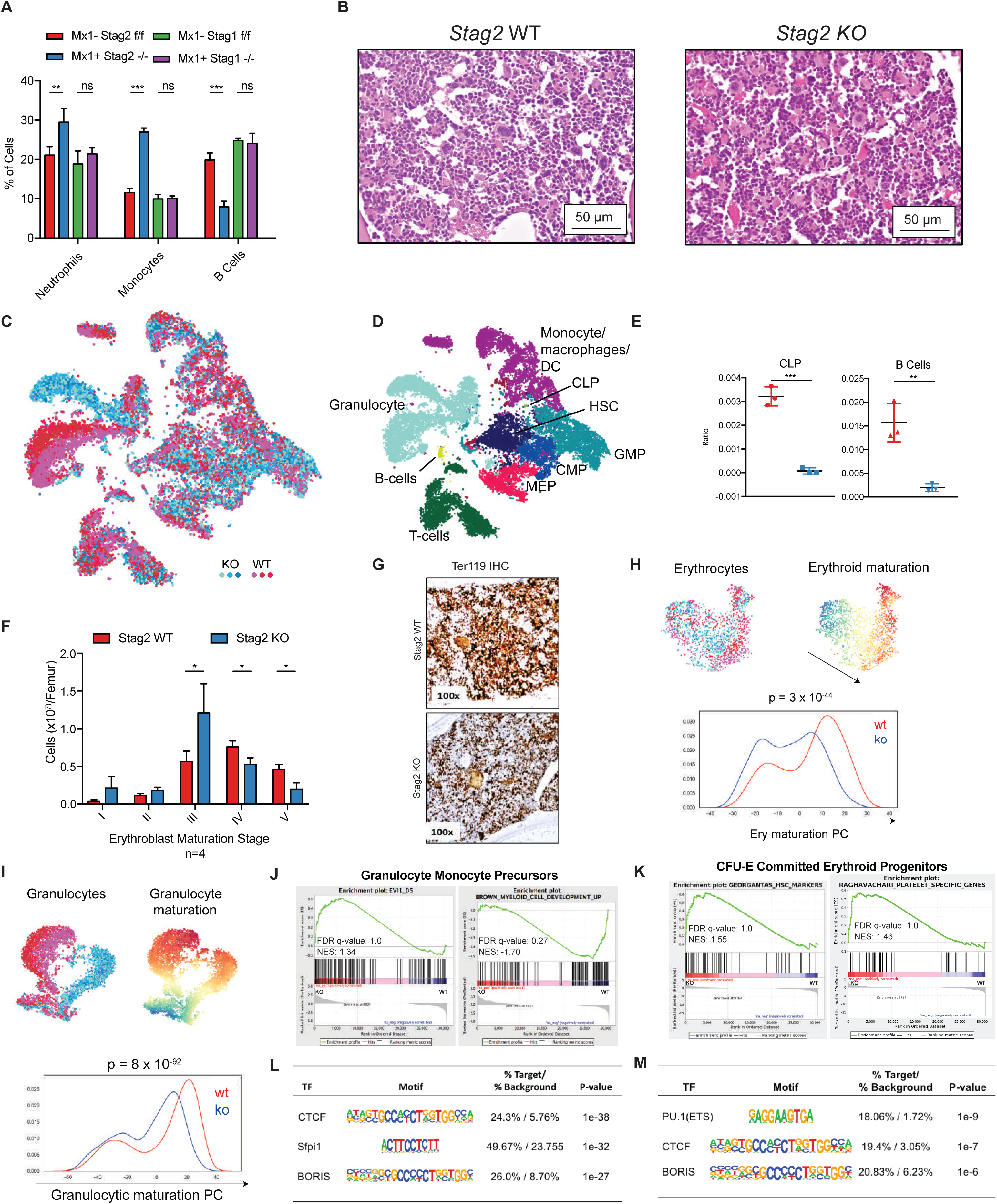
Stag2 alters transcriptional lineage commitment in stem and progenitor cells. **A)** Peripheral blood myeloid skewing with increase in Neutrophils (Gr1^+^Mac1^hi^), Monocytes (Gr1^+^Mac1^lo^), and decrease in B cells (B220^+^) in Stag2 KO mice. No statistical differences were observed in Stag1 KO mice. **B)** Hematoxylin and eosin (H&E) staining of sternal bone marrow from Stag2 KO mice show small, hypolobated megakaryocytes, increased immature myeloid cells, and nuclear-to-cytoplasmic dyssynchrony in erythropoiesis. Single cell RNA sequencing was performed on Stag2 WT (n=3; shades of red) and Stag2 KO (n=3; shades of blue) Lin^−^ HSPC. **C**) t-SNE projection of library-size normalized and log transformed data for complete collection (24,153 cells). Each dot represents a single cell colored by genetic condition (Stag2 KO: shades of blue, Stag2 WT: shades of red). **D**) t-SNE map colored by inferred cell type, as detailed in Supplemental Figure 4. **E**) Frequency of CLPs (left) and B-cells (right) in Stag2 WT (red) and KO (blue) samples; asterisks indicate statistical significance (student’s t test, **p<0.01, ***p<0.001) **F)** Flow cytometric quantification of erythroblast stage in Stag2 WT and KO bone marrow reveals increased erythroblast stage III (p=0.05), but reduction of mature erythroblast stage IV (p=0.02) and stage V (p=0.01). **G)** Immunohistochemical analysis of Stag2 WT and KO bone marrow reveals a marked reduction in Ter-119 expression in Stag2 KO. **H**) Top left: t-SNE projection of library-size normalized and log transformed data for inferred MEP subset (1787 cells). Each dot represents a single cell colored by genetic condition (Stag2 KO: blues, Stag2 WT: reds). Top right: t-SNE projection colored by the second principal component most correlated with erythroid maturation. Bottom: Distribution of cells along erythroid maturation component. **I**) Top left: t-SNE projection of library-size normalized and log transformed data for inferred granulocytic subset (6316 cells). Each dot represents a single cell colored by genetic condition (Stag2 KO: blues, Stag2 WT: reds). Top right: t-SNE projection colored by the first principal component most correlated with granulocyte maturation. Bottom: Distribution of cells along granulocyte maturation component. **J)** Gene-set enrichment analysis of GMP RNAseq shows increased expression of genes in the Evi1 gene set and decreased expression of myeloid development genes. **K)** CFU-E RNAseq shows enrichment for the Georgantas HSC markers gene set and retention of platelet specific genes. Integration of differentially expressed genes in **L)** GMP and **M)** CFU-E populations with ATAC-sequencing commonly have genes with CTCF/CTCFL (BORIS) motif signatures and PU.1 (ETS) motif signatures with decreased expression and decreased accessibility in both populations.

To determine at which level of differentiation multi-lineage dysplasia emerges, we analyzed lineage composition through analysis of scRNAseq data (Figure 2C-D). Supporting the reduction in B cell output observed in the peripheral blood, analysis of scRNAseq data suggested a significant decrease in B cells and in B cell progenitors (Figure 2E). Although the paucity of lymphoid cells precluded further transcriptional tracing, we observed population shifts in erythroid and granulocyte development consistent with observed histologic myelodysplasia. Flow cytometric analysis of erythroblast maturation revealed attenuation at Stage III of erythropoiesis(*34*) (Figure 2F, S6B-C), a stage that is demarcated by condensed cell size as measured by forward scatter (FSC) and decreased CD44 expression relative to prior stages of erythroid differentiation. *Stag2*-deficient erythroid cells were also characterized by morphologic dysplasia; immunohistochemical analysis demonstrated a marked decrease in the proportion of Ter-119^+^ erythroid cells (Figure 2G). By comparison, *Stag1* KO mice did not manifest hematopoietic alterations, morphologic abnormalities, or defects in erythroid differentiation (Figure S6A-C). Transcriptional analysis of the erythroid lineage in *Stag2* deficient cells further supported impaired terminal maturation (p<10^−44^; Figure 2H). Genes defining this principle component define the progression from common myeloid progenitor to megakaryocyte erythroid progenitor to erythroid A (Figure S6D-E). Overall, the most significant transcriptional difference identified in the scRNAseq was impaired granulocytic terminal maturation (p<10^−92^; Figure 2I). Genes defining this principle component define the progression from common myeloid progenitor to granulocyte macrophage progenitor to granulocyte (Figure S6F-G).

Given the alterations in transcriptional output and differentiation potential, we hypothesized that *Stag2* loss led to altered chromatin accessibility at differentially expressed loci. We therefore performed RNA sequencing and ATAC-sequencing on sorted *Stag2* WT and KO GMP and erythroid progenitor (CFU-E: Lineage^−^cKit^+^Sca1^−^Cd34^−^Fcγ^−^Cd105^+^) cells. Compared to WT GMPs, *Stag2-*deficient GMPs had enrichment for a stem-cell gene expression signature associated with increased *Evi1* expression and negative enrichment of myeloid cell development signatures by GSEA (Figure 2J). Similarly, *Stag2-*deficient CFU-Es had increased expression of genes normally expressed in HSCs and in the megakaryocyte lineage, consistent with the increased self-renewal and impaired differentiation seen with Stag2 loss (Figure 2K). Integrated analysis of RNA-seq and ATAC-seq revealed that a significant subset of the downregulated genes also had reduced chromatin accessibility. Both GMPs and CFU-E shared loss of accessibility and expression for genes with either CTCF/CTCFL (BORIS) or PU.1(ETS) motifs (Figure 2L-M). CTCF is a well-described cohesin binding partner(*35*) that cooperates with cohesin to mediate enhancer/promoter associations(*12*). Notably, PU.1 and its target genes have a critical role in determining hematopoietic lineage commitment(*36, 37*).

### Stag1/2 possess shared and independent chromatin binding

It has been shown using *in vitro* model systems that cohesin is essential for genomic cis-interactions and that rapid depletion of cohesin ring members result in complete loss of interphase architecture(*3, 14, 38*). However, these model systems do not reflect a physiologic loss/partial loss of a cohesin complex member akin to the genomic alterations observed in either human cancers or the germline cohesinopathy, Cornelia de Lange syndrome(*39*). As the complete loss of cohesin structural loop components may not be compatible with a viable cell division event, the absence of transcriptional changes observed using *in vitro* systems of cohesin degradation(*14*) may not reflect the pathophysiology of Stag2 loss of function. We next sought to determine how *Stag2*-loss affected chromatin topology in primary cells using Hi-C chromosome conformation capture in hematopoietic progenitor cells. *Stag2* KO cells did not show alterations in topologically associating domain (TAD) boundaries, in global insulation, and did not show differences in the spatially segregated genomic regions known as A/B compartments comparted to WT hematopoietic progenitor cells(*40*) (Figure 3A-C).

**Fig 3.**
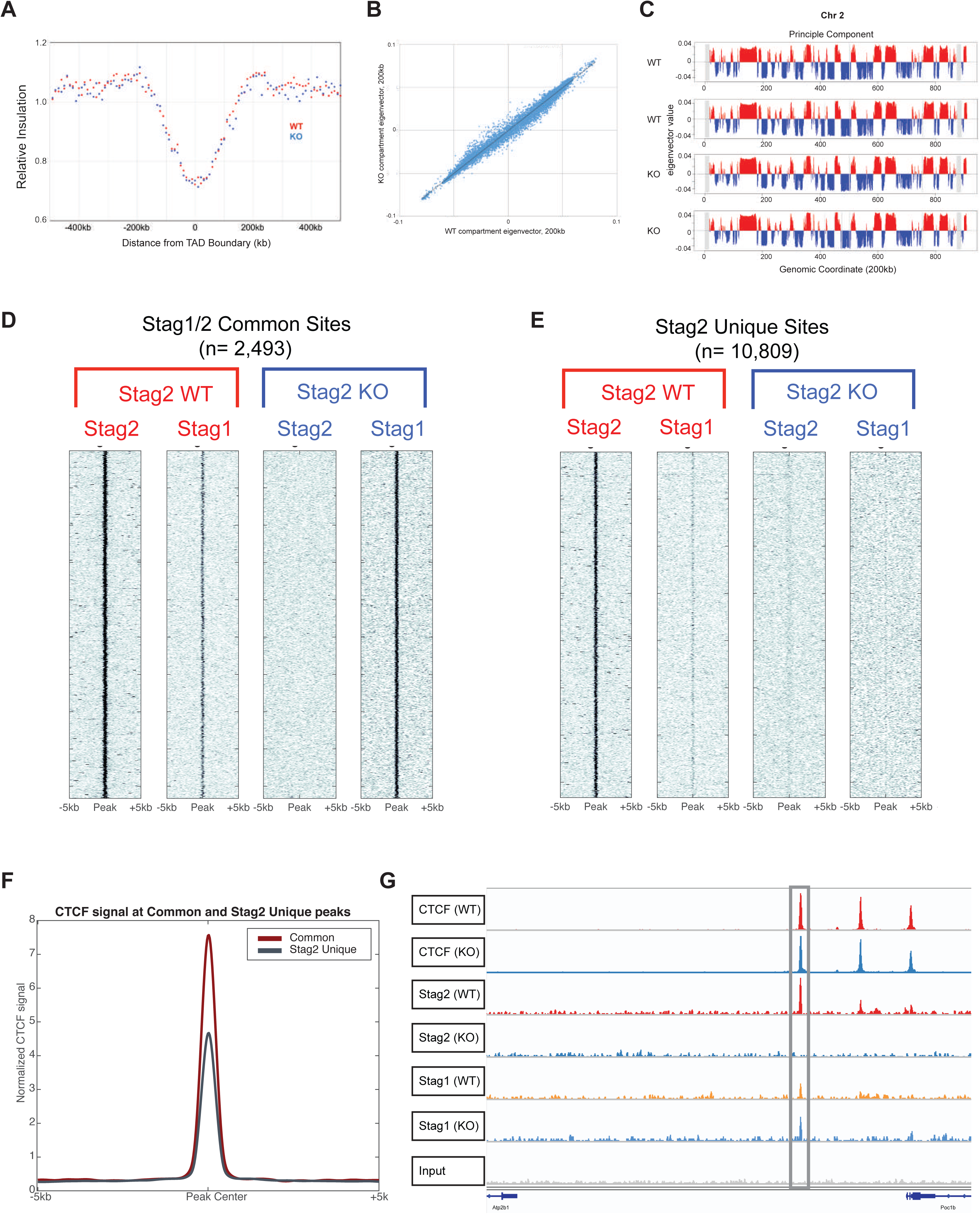
Stag2 and Stag1 possess shared and independent chromatin binding. **A)** Insulation pileup plot from each dot shows the median of the interactions for each 200kb bins located 500kb upstream or downstream of a TAD border, normalized by the median interactions for WT and KO, respectively. **B)** Hi-C insulation scores were normalized by the median interactions for WT and KO, respectively. The diagonal bins were excluded for calculating the median. **C)** A/B compartments of chr2 for each biological replicate. The A compartments, positive PC1 signals, are highlighted in red, while the B compartments, negative PC1 signals, are highlighted in blue. **D)** Chromatin immunoprecipitation and sequencing for Stag1 and Stag2 in Stag2 WT and KO HSPC show a discrete subset of the genome where Stag1 is able to bind Stag2 loci in its absence (common sites) and **E)** discrete loci where Stag1 is unable to bind in the absence of Stag2 (Stag2-specific sites). **F)** Enrichment of Stag1/2-common sites and Stag2-specific sites for CTCF shows strong enrichment in both sets. **G)** IGV track at TAD boundary as measured by Hi-C insulation on chromosome 10 showing CTCF occupancy as well as Stag2 in WT with Stag1 occupancy increased in Stag2 KO HSPC.

Given the lack of changes in global structure, we sought to determine if Stag2 loss would affect the ability of Ctcf of the remaining structural cohesin components Smc1a and Smc3 to localize to canonical cohesion/CTCF binding sites. Chromatin immunoprecipitation sequencing revealed that the occupancy of these three proteins was largely unaffected by the presence or absence of Stag2 in HSPCs (Figure S7A-C). We hypothesized that complete loss of Stag2 might not alter higher order DNA topology due to redundancy with Stag1 at CTCF bound sites. We therefore performed ChIP-seq to delineate Stag2 and Stag1 occupancy in HSPCs from *Stag2* WT and KO mice. We identified a set of peaks where Stag2 and Stag1 were bound in WT mice and where Stag1 occupancy increased in the absence of Stag2 (n=2,493; Figure 3D, S7D-E). Conversely, we identified a second set of peaks where Stag2, but not Stag1, was bound in WT HSPCs and lost in *Stag2* KO cells without compensatory Stag1 occupancy (n=10,809; Figure 3E, S7F-G). Previous work has demonstrated CTCF-dependent and independent localization of the cohesin complex(*41, 42*) which in epithelial cell lines have preferential binding sites for Stag1 and Stag2(*43*). We hypothesized that altered regulation of Stag2-bound loci in WT cells which are not bound by Stag1 in WT or KO cells, were key to the altered differentiation and self-renewal seen in Stag2 KO mice. We found that loci with both Stag2 and Stag1 binding had the highest enrichment for CTCF binding (Figure 3E-F).

### Loss of chromatin insulation impairs transcriptional output

Stag1 was able to bind a subset of loci previously occupied by Stag2 or both Stag1 and Stag2 in WT cells; this included TAD boundaries determined by Hi-C (Figure 3F). By contrast, when we focused on Stag2-only bound sites, we observed locus-specific alterations in short-range interactions (as calculation by the insulation metric; Figure S6B). We quantified these alterations through insulation scores (IS) as a measure of cumulative local contacts and insulation score changes (ISC) that reflect local chromosomal organization changes. To examine whether loss of Stag2 binding affects local chromosomal organization, we compared ISC in WT and *Stag2* KO HSPCs and identified 71,815 ISC bins that showed local chromosomal contact changes (10kb bins with insulation score change less than −0.1 or greater than 0.1, Figure S8A, left panel). We then overlapped ISC bins with Stag1/2 co-bound and Stag2-specific peaks. Annotation analysis of both Stag1/Stag2 common and Stag2-specific peaks showed enrichment of Stag2-specific ISC peaks within promoter regions (Figure S8A, right panel), suggesting that ISC seen in *Stag2-* deficient cells contribute to altered gene expression.

These data suggested that the role of Stag2 in gene regulation and in hematopoiesis might be due to regulation of loci bound by factors other than CTCF. To determine the impact of *Stag2* loss on chromatin structure and transcriptional output we performed RNA-seq and ATAC-seq in WT and Stag2 KO HSPCs. Similar to gene expression/ATAC-seq data in sorted hematopoietic subsets, the majority of genes with significantly altered chromatin accessibility and gene expression in the absence of *Stag2* had decreased accessibility and reduced transcription (Figure 4A). Several of these peaks were in proximity to critical B-cell and myeloid lineage defining transcription factors, including *Ebf1, Pax5*, and *Cebpb*. To uncover transcription factor networks underlying these changes, we performed motif enrichment on peaks with concomitant reduction in accessibility/gene expression. We observed enrichment for Gata, Ebf, and E2A targets (Figure 4B), but not for Cebpb motifs. The genes associated with loss of insulation by ISC significantly overlapped with those loci with decreased accessibility and decreased expression, with *Ebf1* and *Pax5* among the genes with the greatest magnitude of insulation change (Figure 4C, S8B-D). We therefore performed motif analysis on Stag1/2 common and Stag2-specific sites by dichotomizing those that were bound and not bound by CTCF. As expected, loci with shared Stag1/Stag2 occupancy and CTCF binding sites were enriched for CTCF and CTCFL (BORIS) binding motifs. By contrast, Stag2-specific loci (not bound by Stag1) were enriched for key lineage-specific transcription factor motifs including ERG, IRF8, and PU.1 (Figure 4D). These transcription factors are essential for myeloid and lymphoid development (*44, 45*) and attenuated function of these transcription factors has transformative potential in murine(*37*) and human hematopoietic cells(*46-48*).

**Fig 4.**
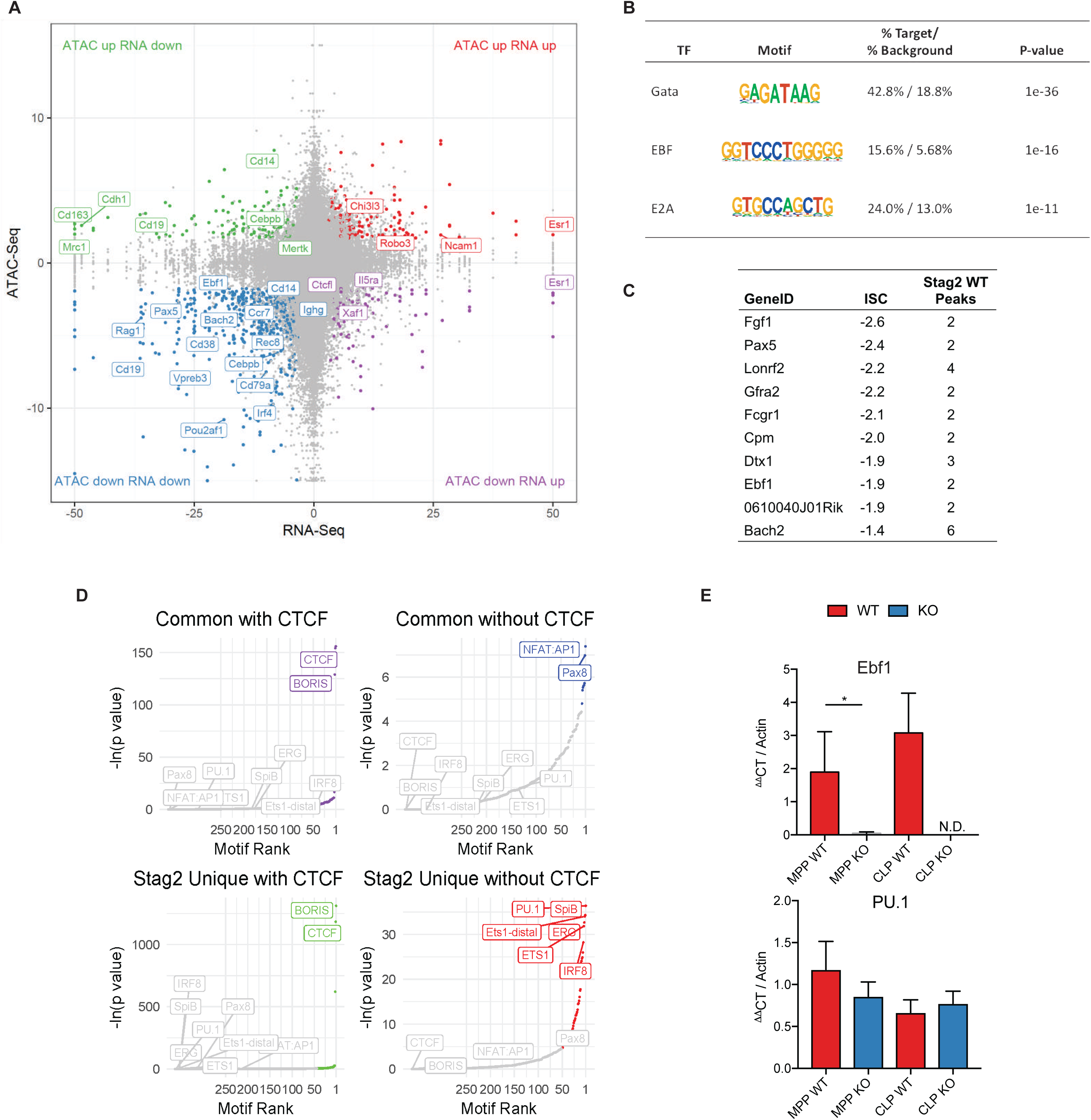
Stag2 loss decreases chromatin insulation and results in impaired transcriptional output. **A)** Intersection of RNAseq and ATACseq in lineage negative bone marrow plotted at Log_2_ fold change of Stag2 KO compared to WT and each point representing a one ATAC peak. The majority of differentially expressed genes lose accessibility, including distinct B-cell regulators (e.g. Ebf1, Pax5, Cd19). **B)** HOMER motif analysis of genes in lower left quadrant of Figure 4A from intersection of RNAseq and ATACseq in lineage negative bone marrow. Genes that are downregulated and lose accessibility are targets of Gata (p=10^−36^), EBF (p=10^−16^), and E2A (p=10^− 11^). **C)** Table of top 10 genes with greatest magnitude of ISC and number of WT Stag2 peaks in the gene. **D)** Motif analyses of common and Stag2-specific sites with and without CTCF show common enrichment for CTCF and CTCFL (BORIS), only Stag2-unique sites bind targets of key lineage priming factors PU.1, SpiB, ERG, ETS1, and IRF8, which Stag1 is unable to bind. **E)** Putative expressers of PU.1 andEbf1 from Stag2 WT and KO bone marrow were sorted and *Ebf1* and *Pu.1* expression was measured by RT-PCR in multipotent progenitors (MPP, p=0.04) and common lymphoid progenitors (CLP, no mRNA detectable in KO). Ebf1 expression was decreased in both populations (MPP p=0.04; CLP not detectable). Pu.1 expression was not statistically different in either population (MPP p=0.13; CLP p=0.32).

### Altered Ebf1 chromatin structure results in B-cell blockade

We next sought to determine if the altered expression of hematopoietic transcription factors was due to alterations in the population distribution in HSPC or was due to altered expression of these factors in purified hematopoietic populations, including in multipotent progenitors antecedent to full lineage commitment. We measured *PU.1* and *Ebf1* expression and B cell differentiation in MPPs and in common lymphoid progenitors (CLP: LSK^+^Cd150^−^Cd48^+^ Cd127^+^) from *Stag2* KO and WT mice. We found that *Ebf1* expression was reduced/absent in purified *Stag2* KO MPPs (p<0.04, Log_2_FC=-1.8) and CLPs (not detectible in KO), while there was no significant difference in *PU.1* expression in either cell type (Figure 4E). Consistent with these data, B cell maturation was blocked from the pro-B cell to early pre-B cell stage(*49*) in *Stag2* KO mice, with almost no late pre-B cells (Figure 5A). These findings were further bolstered by *in vitro* colony assays in B cell cytokine enriched methylcellulose where *Stag2* KO HSPCs failed to generate B cell colonies (Figure 5B). We next investigated if the same was true in Stag2-mutant MDS patients by enumerating B cells and B cell progenitors in STAG2-mutant MDS patient bone marrow samples compared to a cohort of non-MDS individuals with cytopenias (Figure 5C, S8E). Immature B cells (CD34+CD19+) were decreased by 1.7-fold (p<0.01) in STAG2-mutant MDS patients, demonstrating that altered B-cell lineage commitment is observed in STAG2-mutant MDS.

**Fig 5.**
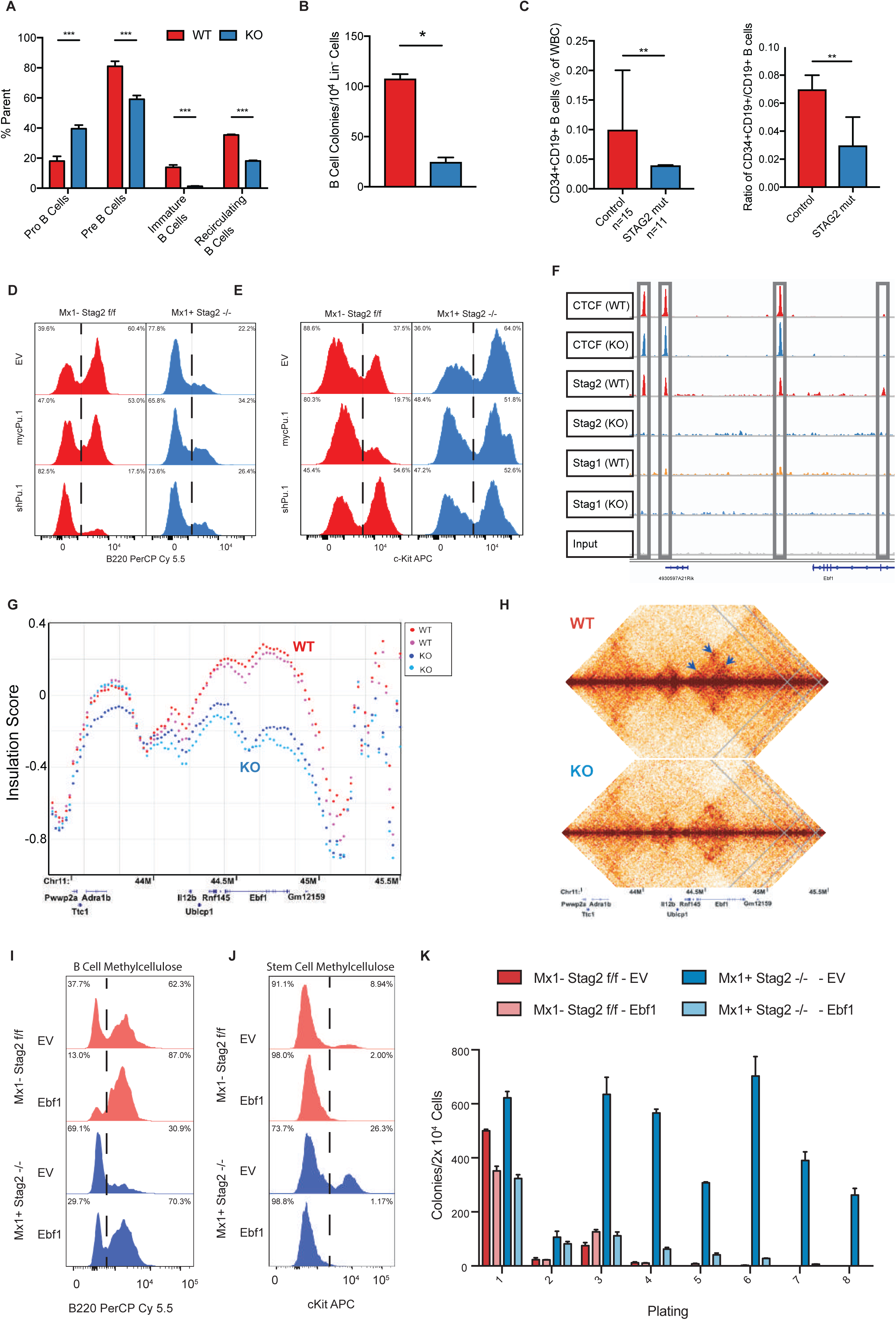
Altered Ebf1 chromatin structure results in a blockade of B cell development. **A)** Enumeration of B lymphocyte stages of maturation in Stag2 WT and KO mice. Bone marrow was analyzed using flow cytometry for pro-(Cd43^+^) and pre-(Cd43^−^) B-cells in parent gate B220^lo^IgM^−^ showing B-cell development was impeded in the pro-B to pre-B cell stage (p=0.003). Immature (B220^lo^IgM^+^) and recirculating B cells (B220^hi^IgM^+^) were analyzed as a percentage of live singlets, which were both markedly reduced in Stag2 KO mice (p<0.001 for both populations). **B**) Methylcellulose colony assay using IL-7, SCF, and Flt3-L enriched media shows reduction in the number of B-cell colonies able to be formed by Stag2 KO bone marrow compared to WT (p=0.003). **C)** Enumeration of immature B cells (CD34+CD19+) and ratio compared to mature B cells (CD34-CD19+) show reduced immature B cells and immature:mature ratio in STAG2 mutated MDS patients (n=11) compared to controls (n=15). Stag2 WT and KO HSPC were infected with lentivirus containing GFP-tagged empty vector, GFP-mycPu.1, or GFP-shPu.1; and GFP^+^ cells were plated in either B cell colony methylcellulose or stem cell methylcellulose. Cells plated in methylcellulose were harvested after 7 days and analyzed by flow cytometry for **D)** mature cell markers including the B marker B220 or **E)** stem cell markers including cKit. **F)** IGV track of the Ebf1 locus with Stag2 and Ctcf binding at 4 distinct sites which are lost in Stag2 KO and not bound by Stag1 either in WT or KO. **G)** ISC plotted across Ebf1 for Stag2 WT (n=2; shades of red) and KO (n=2; shades of blue) shows marked loss of insulation. **H)** Contact map of Ebf1 shows loss of cis-interaction at three loci (arrows). **I)** Lineage negative Stag2 WT and KO marrow were infected with retrovirus containing GFP-tagged empty vector or GFP-pCMV-Ebf1, and GFP^+^ cells were plated in either B-cell colony methylcellulose or stem cell methylcellulose. As in C-D, cells were harvested after 7 days and analyzed by flow cytometry for **I)** mature cell markers including the B marker B220 or **J)** stem cell markers including cKit. **K)** Cells plated in stem cell methylcellulose were serially replated with exhaustion of Stag2 KO cell overexpressing Ebf1.

Given that PU.1 can regulate myeloid and lymphoid lineage commitment(*37*) and the enrichment for PU.1 motif/targets in our epigenomic/transcriptional analysis, we investigated the role of PU.1 in Stag2-deficient alterations in B-cell lineage commitment and self-renewal. PU.1 silencing reduced B-lineage output derived from WT hematopoietic cells, similar to the effects of Stag2 deletion (Figure 5D). However, the reduced B-cell lineage output of *Stag2* KO cells was not further attenuated by PU.1 silencing as assessed by flow cytometric analysis of B220 expression and B-lineage colony formation (Figure 5D). By contrast, overexpression of PU.1 was not sufficient to restore B220 expression in B cell methylcellulose cultures (Figure 5D) or to attenuate the increased expression of the stem/progenitor marker cKit seen on *Stag2-*deficient hematopoietic cells (Figure 5E). Moreover, overexpression of PU.1 failed to attenuate the capacity of *Stag2-* deficient hematopoietic cells to serially replate (Figure S8F-H).

This led us to hypothesize that key transcription factors downstream of PU.1 which are bound by Stag have locus specific alterations in conformation in the absence of Stag2 that lead to alterations in target gene expression with functional import. *Ebf1* has four Stag2 bound sites in hematopoietic progenitors, all of which are not bound by Stag1 in WT or KO cells (Figure 5F). As noted above, we observed near-complete abrogation of *Ebf1* transcription from the onset of B-lineage commitment (Figure 4E). We observed a significant reduction in local insulation throughout the *Ebf1* locus (Figure 5G) including loss of cis-interactions at three Stag2 binding sites (Figure 5H). Consistent with this hypothesis, expression of *Ebf1* in Stag2 KO progenitors (Figure S8I) fully restored B-lineage marker expression and B-cell colony formation *in vitro* (Figure 5I). Moreover, *Ebf1*-expressing Stag2 KO progenitors had decreased cKit expression (Figure 5J) and *Ebf1* restoration abrogated the serial replating capacity of *Stag2*-deficient cells (Figure 5K). Taken together, these data demonstrate a key role for Stag2-mediated regulation of Ebf1 expression in modulating the balance between hematopoietic differentiation and self-renewal.

## Discussion

The cohesin complex is essential in pleiotropic cellular and gene regulatory functions, including in chromosome segregation(*10, 11*) and in maintaining DNA interactions(*3, 14*) which regulate gene expression(*13, 15, 16*). Here we demonstrate a non-redundant role of Stag2 and Stag1 in hematopoiesis, with a specific role for Stag2 in regulating the balance between hematopoietic differentiation and self-renewal. In addition, we show that Stag2 bound loci that are not bound by Stag1 undergo locus specific changes in local interactions that, in the absence of Stag2, leads to altered expression of key loci governing hematopoietic function. Taken together, these data illustrate a critical link between Stag2, DNA interactions, and gene regulation that contributes to hematopoietic transformation.

Stag2 and Stag1 have both shared and distinct roles(*43*). These paralogs have a redundant role in chromatid separation such the presence of either protein within the cohesin complex is adequate for intact sister chromatid alignment. In contrast to deletion of core cohesin ring components, which abrogates the survival of hematopoietic cells(*19*), somatic deletion of either *Stag1* or *Stag2* was compatible with hematopoietic stem cell viability, whereas loss of both genes led to hematopoietic failure and defective chromosomal segregation. Our *in vivo* studies underscore previous *in vitro* dependency studies(*20*) and suggest that therapies which directly or indirectly alter Stag1 function will induce a specific dependency in Stag2 mutant cancer cells. However, increased self-renewal, impaired lineage commitment, and progressive hematopoietic alterations were observed with Stag2 deletion but not with Stag1 loss. These studies illustrate that Stag2 has a key role in hematopoiesis and in hematopoietic transformation that is not shared by Stag1. Future studies will illustrate how Stag2 loss of function cooperates with other alleles to promote leukemic transformation and how co-occurring disease alleles cooperate with the chromatin architecture alterations induced by *Stag2* loss.

Our data suggest that the role of Stag2 in regulating hematopoiesis and in hematopoietic transformation is Stag1-independent. Key hematopoietic regulators which show differential expression, accessibility, and insulation in the absence of Stag2 are bound by Stag2 and not by Stag1. This includes several key genes essential for lineage commitment to lymphoid fate, including the PU.1 targets *Ebf1* and *Pax5*. Previous studies demonstrated that PU.1-mediated regulation of *Ebf1* is a key event in B-cell lineage commitment(*50, 51*), and PU.1 has been defined to be a pioneer factor with nucleosome remodeling capacity. Given these data and the known role of Ebf1 in B cell commitment/differentiation (*22*), we show that PU.1 restoration cannot rescue the impact of Stag2 loss on hematopoietic self-renewal and lineage commitment without the permissive chromatin context regulated by Stag2. By contrast, restoration of Ebf1 expression restores differentiation and abrogates the aberrant self-renewal of Stag2 deficient cells.

Our data illustrate a key role for Stag2 in the balance between self-renewal and differentiation, a critical feature of transformation across different tumor types. Moreover, the observation that reactivation of the Stag2-target Ebf1 can reverse these alterations suggests a key role for Stag2-regulated transcription factor networks in tumor suppression, which is reversible when key networks are restored in Stag2 mutant cells. We hypothesize that Stag2 loss-of-function induces lineage specific alterations in transcription factor expression and function in a spectrum of malignant contexts. Most importantly, our studies identify a key pathophysiologic role for mutations in the factors that govern gene regulatory architecture in malignant transformation. Our data point to the reversibility of transcriptional dysregulation which mediates aberrant self-renewal and differentiation in cancer cells and suggest that therapies that reverse these alterations may have therapeutic benefit in cohesin-mutant cancers.

## Acknowledgments

We acknowledge the use of the Integrated Genomics Operation Core, funded by the Memorial Sloan Kettering Cancer Center Support Grant NIH P30 CA008748, and the Marie-Josée and Henry R. Kravis Center for Molecular Oncology. We would like to thank Dr. Gauri Nandakumar and the staff of the Cytogenetics Core Facility for their assistance in imaging and processing of cytogenetic data.

## Funding

This work was supported by a National Cancer Institute R35 CA197594-01A1 (R.L.L.), National Cancer Institute R01 CA216421 (R.L.L.), National Cancer Institute PS-OC U54 CA143869-05 (R.L.L. and J.D.), and Leukemia & Lymphoma Society Translational Research Foundation 6499-17 (R.L.L.). A.D.V. is supported by a National Cancer Institute career development grant K08 CA215317, the William Raveis Charitable Fund Fellowship of the Damon Runyon Cancer Research Foundation (DRG 117-15), and an EvansMDS Young Investigator grant from the Edward P. Evans Foundation. R.L.B. is supported by the Sohn Foundation Fellowship of the Damon Runyon Cancer Research Foundation (DRG 22-17). T.B. is supported by the William C. and Joyce C. O’Neil Charitable Trust, Memorial Sloan Kettering Single Cell Sequencing Initiative. J.D. is an investigator of the Howard Hughes Medical Institute.

## Author Contributions

Conceptualization, A.D.V., R.L.B., R.L.L. Methodology, A.D.V., R.L.B., Y.L., V.L., W.X., B.H.D., S.E.E., A.N., M.W., T.B., E.A., S.L., C.J.O., D.P., J.D., R.K., R.L.L. Investigation, A.D.V., R.L.B., Y.L., W.X., B.H.D., S.E.E., A.N., J.P., S.B., M.W., T.B., S.L., C.J.O., J.D., R.K., R.L.L. Writing – Original Draft, A.D.V., R.L.B. Writing – Review & Editing, A.D.V., R.L.B., Y.L., B.H.D., S.E.E., A.N., M.W., T.B., S.L., C.J.O., J.D., R.K., R.L.L. Funding Acquisition, R.L.L. Resources, J.D. R.L.L. Supervision, R.L.L.

## Competing interests

A.D.V. received travel support from Mission Bio and is on the Editorial Advisory Board of Hematology News. J.D. is on the Scientific Advisory Board of Arima Genomics, and a consultant for Omega Therapeutics. R.L.L. is on the supervisory board of Qiagen and is a scientific advisor to Loxo, Imago, C4 Therapeutics and Isoplexis. He receives research support from and consulted for Celgene and Roche and has consulted for Janssen, Astellas, Morphosys and Novartis. He has received honoraria from Roche, Lilly and Amgen for invited lectures and from Gilead for grant reviews. R.L.B, Y.L., B.H.D., S.E.E., A.N., M.W., T.B., C.J.O., and R.K. have nothing to disclose.

## Supplementary Materials

Materials and Methods

Figures S1-S8

