## Supplemental Figures for "Stag1 and Stag2 regulate cell fate decisions in hematopoiesis through non-redundant topological control"

**This PDF file includes:**

Materials and Methods
Figs. S1 to S8
Captions for Data S1 to S8

#### Materials and Methods

##### Animals.

All animals were housed at Memorial Sloan Kettering Cancer Center. All animal procedures were conducted in accordance with the Guidelines for the Care and Use of Laboratory Animals and were approved by the Institutional Animal Care and Use Committees at Memorial Sloan Kettering Cancer Center.

##### Generation of Stag1/2-deficient mice.

The *Stag2* and *Stag1* conditional allele were each deleted by targeting exon 7. Two *LoxP* sites flanking exon 7 and an *Frt*-flanked neomycin selection cassette were inserted in the upstream intron (Figure S1A). 10 µg of the targeting vector was linearized by NotI and then transfected by electroporation of BAC-BA1 (C57BL/6 × 129/SvEv) hybrid ES cells. After selection with G418 antibiotic, surviving clones were expanded for PCR analysis to identify recombinant ES clones. Secondary confirmation of positive clones identified by PCR was performed by Southern blotting analysis. DNA was digested with BamHI and electrophoretically separated on a 0.8% agarose gel. After transfer to a nylon membrane, the digested DNA was hybridized with a probe targeted against the 3′ or 5′ external region. DNA from C57BL/6 (B6), 129/SvEv (129), and BA1 (C57BL/6 × 129/SvEv; Hybrid) mouse strains was used as WT controls. Positive ES clones were expanded and injected into blastocysts. The generated mice (*Stag2*$^{fl/fl}$ and *Stag1*$^{fl/fl}$) were initially crossed to a germline *Flp*-deleter (The Jackson Laboratory), to eliminate the neomycin cassette, and subsequently to the IFN-α–inducible *Mx1-cre* (The Jackson Laboratory) (*52, 53*). Mice were backcrossed for six generations to C57BL/6 mice. *Stag2*$^{fl/fl\ or\ y}$, *Stag2*$^{fl/+\ or\ y}$, and *Stag2*$^{+/+\ or\ y}$ littermate mice were genotyped by PCR with primers Stag2-NF (5′-CACTCATGCTGGCAAGTATTGTAC-3′) and Stag2-NR (5′-AACAGCCTGAGCAAAGAATCCAAAG-3′) and Stag2-3' (5′-TGTGTGCCTCTTTGAACAATGCCC-3′) using the following parameters: 94°C for 3 min, followed by 35 cycles of 94°C for 15 s, 64°C for 30 s, and 72°C for 90 s, and then 72°C for 5 min. The WT allele was detected as a band at 385 bp, whereas the floxed allele was detected as a band of 554 bp. Excision after Cre recombination was confirmed by PCR with primers to detect a band at 294 bp. *Stag1*$^{fl/fl}$, *Stag1*$^{fl/+}$, and *Stag1*$^{+/+}$ littermate mice were genotyped by PCR with primers Stag1-CommonF (5′-GACTGGTATCTGACGGCTTATACC-3′) and Stag1-CommonR (5′-CACTGAGGACCAGGCATTGTAAGG-3′) and Stag1-FloxR (5′-TGAACTGATGGCGAGCTCAGACC-3′) using the following parameters: 94°C for 3 min, followed by 35 cycles of 94°C for 15 s, 62°C for 30 s, and 72°C for 90 s, and then 72°C for 5 min. The WT allele was detected as a band at 1104 bp, whereas the floxed allele was detected as a band of 745 bp. Excision after Cre recombination was confirmed by PCR with primers to detect a band at 619 bp. The following mouse lines have been donated to The Jackson Lab and will be available: *Stag2*$^{flox}$ (JAX#030902); *Stag1*$^{flox}$ (JAX#030904).

**In vivo experiments.**
*Mx1-cre*$^{+}$ *Stag2*$^{fl/fl}$ conditional KO and Cre$^{-}$ *Stag2*$^{fl/fl}$ control WT mice received four intraperitoneal injections of polyinositric:polycytidinic acid (PIpC) every other day at a dose of 20 mg/kg of body weight starting at 6 weeks after birth. Mice were analyzed between 3 and 60 weeks of age. BM, spleen, and peripheral blood were analyzed by flow cytometry. Formalin-fixed paraffin-embedded tissue sections were stained with hematoxylin and eosin (H&E). Peripheral blood was smeared on a slide and stained using the Wright-Giemsa staining method. Tissue sections and blood smears were evaluated by a hematopathologist (B. Durham). Deletion of the *Stag1 and Stag2* allele and transcript was measured by genomic PCR, qRT-PCR, and Western blot analysis. All experiments were repeated and results confirmed in male *Mx1-cre*$^{+}$ *Stag2*$^{fl/y}$ conditional KO and Cre$^{-}$ *Stag2*$^{fl/y}$ control WT mice.

**BM transplantation.**
Freshly dissected femurs and tibias were isolated from *Stag1*$^{fl/fl}$ and *Stag2*$^{fl/fl}$ CD45.2$^{+}$ or *Mx1-cre*$^{+}$ *Stag1*$^{fl/fl}$ and *Stag2*$^{fl/fl}$ CD45.2$^{+}$ mice. Bones were transected at the epiphyses and were centrifuged at 4°C to extract whole bone marrow into PBS and RBCs were lysed in ammonium chloride-potassium bicarbonate lysis buffer for 10 min. After centrifugation, cells were resuspended in PBS plus 3% FBS, passed through a cell strainer, and counted. Finally, 0.5 × $10^6$ total BM cells from *Stag1*$^{fl/fl}$ and *Stag2*$^{fl/fl}$ CD45.2$^{+}$ or *Mx1-cre*$^{+}$ *Stag1*$^{fl/fl}$ and *Stag2*$^{fl/fl}$ CD45.2$^{+}$ mice were mixed with 0.5 × $10^6$ WT CD45.1$^{+}$ support BM and transplanted via tail vein injection into lethally irradiated (two times 450 cGy) CD45.1$^{+}$host mice. Chimerism was measured by FACS in peripheral blood at 4 weeks after transplant (week 0, pre-PIpC). Chimerism was followed via FACS in the peripheral blood every 4 weeks (week 0, 4, 6, 8, 12, and 16 after PIpC injection). Additionally, for each bleeding, whole blood cell counts were measured on a blood analyzer, and peripheral blood smears were scored. Chimerism in the BM and spleen was evaluated at 16 weeks via animal sacrifice and subsequent FACS analysis.

**In vitro colony-forming assays.**
BM of *Stag1*$^{fl/fl}$ and *Stag2*$^{fl/fl}$ and littermate *Mx1-cre Stag1*$^{fl/fl}$ and *Stag2*$^{fl/fl}$ mice were extracted and seeded at a density of 20,000 cells/replicate into cytokine-supplemented methylcellulose medium

(Methocult M3434; STEMCELL Technologies) or methylcellulose medium for mouse pre-B cells (M3630) supplemented with FLT3L (20 ng/mL), SCF (100 ng/mL), and IL7 (10 ng/mL)(*54*). Colonies propagated in culture were scored at day 7. Representative colonies were isolated from the plate for cytospins and flow cytometry. Remaining cells were resuspended and counted, and a portion was taken for replating (20,000 cells/replicate) for a total of eight platings. Cytospins were performed by resuspending in warm PBS and spun onto the slides at 350 *g* for 5 min. Slides were air-dried and stained using the Giemsa-Wright method.

**Antibodies, FACS, and Western blot analysis.**
Antibody staining and FACS analysis was performed as previously described (*55*). BM or spleen mononuclear cells were stained with a lineage cocktail comprised of antibodies targeting CD4, CD8, B220, NK1.1, Gr-1, CD11b, Ter119, and IL-7Rα. Cells were also stained with antibodies against c-Kit, Sca-1, FcγRII/III, and CD34. Cell populations were analyzed using a Fortessa Flow Cytometer (BD) and sorted with a FACS-SH800 instrument (Sony). All FACS antibodies were purchased from BD or eBioscience. We used the following antibodies: c-Kit (2B8), Sca-1 (D7), Mac-1/CD11b (M1/70), Gr-1 (RB6-8C5), NK1.1 (PK136), Ter-119, IL7-Rα (A7R34), IgD (11-26c.2a), IgM (RMM-1), CD34 (RAM34), FcγRII/III (2.4G2), CD4 (RM4-5), CD8 (53-6.7), CD19 (HIB19), CD43 (1B11), CD44 (IM7), CD45.1 (A20), CD45.2 (104), CD45R/B220 (RA3-6B2), CD71 (R17217), CD105 (MJ7/18), CD150 (9D1), and CD48 (HM48-1). The following antibodies were used for Chromatin Immunoprecipitation and Western blot analysis: Stag2 (Bethyl, A302-580A), Stag1 (Bethyl, A302-579A), Smc3(Bethyl, A300-060A), Smc1a (Active Motif, 61067), Ctcf (Cell Signaling, 3418S), and Actin (CalBiochem, JLA-20).

Multiparameter flow cytometry was performed on bone marrow aspirates at diagnosis for patients with STAG2 mutant MDS and matched controls patients with nonmalignant cytopenias. Briefly, up to 1.5 million cells from freshly drawn bone marrow aspirate were stained with a 10-"color" panel (CD15-FITC, CD33-PE, CD117-PC5, CD13-PE-Cy7, CD34-APC, CD71-APC-A700, CD38-APC-A750, HLA-DR-PAC Blue, CD45-V500C, and CD19-BV605), washed, and acquired on a Canto-10 cytometer (BD Biosciences, San Jose, CA). The results were analyzed with custom Woodlist software (generous gift of Wood BL, University of Washington). In order to enumerate the B cells, CD19+CD15-CD33-CD13- cells were gated. Plasma cells were excluded based on the bright CD38 expression. CD34+/CD19+ cells were considered as immature B cells. This study was approved by Institutional Review Board at the MSKCC (protocol #16-1591).

**Histological analyses.**
Mice were sacrificed and autopsied, and dissected tissue samples were fixed for 24 h in 4% paraformaldehyde, dehydrated, and embedded in paraffin. Paraffin blocks were sectioned at 4 µm and stained with H&E. Images were acquired using an Axio Observer A1 microscope (Carl Zeiss).

**Peripheral blood analysis.**
Blood was collected by submandibular bleeding using a 5mm lancet (MEDipoint Inc). Automated peripheral blood counts were obtained using a ProCyte Dx (IDEXX Laboratories) according to standard manufacturer's instruction. Differential blood counts were realized on blood smears stained using Wright-Giemsa staining and visualized using an Axio Observer A1 microscope.

**Cytogenetic Analysis and Metaphase Karyotyping.**
Bone marrow acquired at necropsy was resuspended in cytokine enriched media containing 5mL RPMI 10% FCS supplemented with 2mM L-glutamine and after documented excision or 8 weeks after PIpC. Harvested cells were cultured in T25 tissue culture flasks with 25 uL of Colcemid (10mg/mL)(Gibco Life Technologies, Inc) for 45 minutes and 4 hours respectively, resuspended

in 0.075 mol/L KCl for 10 minutes at 37°C and fixed in methanol-acetic acid (3:1). Metaphases were scored and counted. Chromosome analysis was performed on a minimum of 20 DAPI-banded metaphases and all metaphases were fully karyotyped. Low depth whole genome sequencing was performed to assess genome wide copy number. One microgram of genomic DNA was sheared using the Covaris instrument to +/- 300bps. Fragmented DNA was end-repaired, A-tailed, and ligated to Illumina TruSeq dual indexed adaptors using standard methods. Indexed libraries were enriched by PCR amplified, quantified, pooled and sequenced in multiplex fashion on an Illumina HiSeq instrument to obtain roughly 1 million sequencing reads per sample, sufficient to call copy number variation at a bin resolution of 150kb)(*56*). Data was processed for copy number analysis as described previously(*57*).

**RNA-Seq and quantitative real-time PCR (qRT-PCR) analysis.**
For qRT-PCR experiments, all samples were prepared in biologic triplicate. Whole BM was negatively selected for lineage markers using antibodies conjugated to magnetic beads and separated using EasySep Mouse Hematopoietic Progenitor Cell Isolation Kit (STEMCELL Technologies). Total RNA was isolated using the Trizol (Invitrogen), and cDNA was synthesized using the Verso cDNA Synthesis kit (Fisher). Quantitative PCR was performed using Taqman reagents and probes (Thermo Fisher) for ActinB (Mm02619580_g1), Spi1, and Ebf1 (Mm00432948_m1) and FastStart Universal SYBR Green (Sigma) and primers for Stag2 (F: 5'-TGCTATGCAGTCGGTGGTAG-3') and (R: 5'-AGGACCAGCCATGGTAAGTG-3'), Stag1 (F: 5'-CTACAAGCATGACCGGGACAT-3') and (R: 5'-AGGGTACTTGTATGCCTAAAAGC-3'), and ActinB (F: 5'-GGCTGTATTCCCCTCCATCG-3') and (R: 5'-CCAGTTGGTAACAATGCCATGT-3'). For mRNA-Seq analysis, samples were prepared and analyzed in biologic triplicate. For RNA-sequencing, RNA was isolated by TriZOL extraction from sorted-cell population or lineage negative bone marrow as indicated. RNA-sequencing libraries were generated by 3' sequencing and SMART-Seq2 amplification and sequenced on an Illumina HiSeq 2500. Fastq files were mapped to the mouse genome (mm9) and reads counts per gene were quantified using STAR(*58*) with default parameters and genecode (vM1) annotation file. Differentially expressed genes (DEGs) were identified with DESeq2(*59*), with a fold change cutoff of +/- 2 and a FDR of 10%. Motif enrichment on DEGs was performed with HOMER using a window of -1kb to +100bp around the transcription start site. Gene ontology analysis was also performed using HOMER.

**Single cell RNA-sequencing and data analysis.**
Lineage$^{-}$bone marrow cells from 3 Stag2 WT and 3 Stag2 KO mice were sorted for viability (4′,6-diamidino-2-phenylindole (DAPI)-negative). Individual samples were loaded on 10X Genomics Chromium System aiming to generate 7,000 Gel Beads in Emulsion (GEMs) per sample. scRNA-seq libraries were prepared following 10X Genomics protocols (Chromium Single Cell 3' Reagent Kits User Guide v2 Chemistry). Libraries were sequenced on NovaSeq 6000 (Illumina) system (S2 flow cell, paired-end) recovering a median of 239,350,330 reads/sample. FASTQ files were processed using the Sequence Quality Control (SEQC) pipeline(*60*) and reads were aligned to the mouse genome mm38, resulting in a median of 4226 cells/samples with a median of 5036 molecules/cell. Cells from the lower molecule counts, determined by lower mode of molecule counts distribution (5.8% of cells), were additionally filtered out to remove putative empty droplets, resulting in a final collection of 24,153 cells. The resulting count matrices from all samples were then combined and normalized to median library-size and log transformed shown in Figures 2C-D and S4C. For individual subsets of granulocytes, MEP, HSCs, cells were normalized within the subset. Ribosomal genes were excluded in downstream analyses. In the global cohort, t-SNE was performed on 20 principal components, explaining ~83.6% of data variance, with a perplexity of 150. For subpopulations analyses, the number of PCs considered was determined based on the knee point of explained variance (granulocytes: 8, MEP: 14, HSC: 7) and perplexity was fixed at 150.

In order to annotate principal components (Figure 1G, 2H, 2I), the Pearson correlation between each principal component and expression of gene signatures was computed. Gene lists were sorted by correlation and ranked Gene Set Enrichment Analysis (GSEA) was performed. In addition, expression of most correlated and anticorrelated was assessed in bulk RNA-sequencing data of sorted populations to identity components defined by lineage maturation.

To annotate subsets of cells, we performed clustering of normalized and log transformed data using Phenograph (*61*). Then, Pearson correlation between the centroid of each cluster and standardized bulk RNA-seq data on sorted subsets (from Haemopedia-Mouse RNAseq,(*62*)) was computed (Figure S4D). The subset with highest correlation to each cluster was used to annotate the cluster, as shown in Figures S4D, 2D. The cluster annotations were confirmed through studying differentially expressed genes in each cluster. Differential expression analysis was performed using the Wasserstein distance(*63*) between normalized and log-transformed expression in cells from each inferred lineage and all other cells (Figure S4E).

**PU.1/EBF1 overexpression and knockdown**
Lentiviral constructs expressing PU.1-IRES-GFP, PU.1 shRNA-IRES-GFP, and GFP alone were generously provided by the laboratory of Dr. Ulrich Steidl (*37, 64*). Whole BM was negatively selected for lineage markers using antibodies conjugated to magnetic beads and separated using EasySep Mouse Hematopoietic Progenitor Cell Isolation Kit (STEMCELL Technologies). Lineage depleted BM cells were transduced at a cell density of 5 $x10^5$ using virus concentrated through ultracentrifugation. After 48h cells were sorted for GFP expression. Overexpression/knockdown was confirmed using qRT-PCR. Retroviral constructs expressing EBF1-IRES-GFP and GFP alone were generously provided by the laboratory of Dr. Charles Mulligan. Lineage depleted BM cells were transduced at a cell density of 5 $x10^5$ using virus concentrated using Retro-X concentrator (Clontech). After 48h cells were sorted for GFP expression. Overexpression was confirmed using qRT-PCR.

**ATAC-sequencing**
Chromatin accessibility assays utilizing the bacterial Tn5 transposase were performed as described (*65*) with minor modifications. Cells ($10 \times 10^3$) were lysed and incubated with transposition reaction mix for 30 minutes at 37°C. Samples were PCR-amplified and sequenced on an Illumina NextSeq 500.

**Chromatin Immunoprecipitation-sequencing**
Chromatin immunoprecipitation (ChIP) was performed as previously described. Cells were cross-linked with 1% formaldehyde (Sigma, F1635) at 37°C for 15 min and quenched with 0.125 M glycine for 5 min at room temperature. Cells pellets were obtained by centrifugation with 3000 RPM for 5 minutes at 4C and froze in liquid nitrogen immediately before transferring to the -80C freezer. ~25 millions cells were used for one ChIP reaction. Cell pellets were thawed on ice, resuspended in 1ml SDS lysis buffer (1% SDS, 10mM EDTA, 50mM Tris-HCl pH8) containing proteinase inhibitor and phosphatase inhibitor for one reaction in eppendorf tube and incubated on ice for 10 minutes. Sonication was performed on a Branson Sonifier 250 with a 20% amplitude setting for 5.5 minutes (10 second on/off pulsing). Sonication product were spun down at 14000 RPM at 4C for 10 minutes and 1ml supernatant containing chromatin and DNA were transferred to a falcon tube containing 9ml ChIP dilution buffer (0.01%SDS, 1.1%TritonX-100, 1.2mM EDTA, 16.7mM Tris-HCl pH8, 167mM NaCl) with proteinase and phosphatase inhibitor. 50ul of Dynbeads (Life technologies 10009D) were added to samples and incubated at 4C with rotation for 1 hour. After-pre clearing, dynbeads beads were removed and 200ul of sample were collected

as 2% input separately. 5 µg antibodies were added to the pre-cleared samples for overnight incubation at 4C with rotation. 200ul Dynbeads were added into one ChIP reaction and incubated for 4-6 hours at 4C with rotation. Dynabeads were collected by centrifugation with 3000 RPM at 4C for 5 minutes and washed in 1ml low salt buffer (0.1% SDS, 1% TritonX-100, 2mM EDTA, 20mM Tris-HCl pH8, 150mm NaCl) for 5 minutes at 4C with rotations. Then beads were washed in 1ml high salt buffer (0.1% SDS, 1% TritonX-100, 2mM EDTA, 20mM Tris-HCl pH8, 500mm NaCl) twice and TE buffer (10mM Tris-HCl pH8, 1mM EDTA) twice for 5 minutes at 4C with rotation. After the last wash, beads were resuspended in 250µl elution buffer (1% SDS, 0.1M NaHCO3) and incubated in a thermomixer (850RPM) for 15 minutes at 60C. Supernatant were collected and added with 5M NaCl for overnight decrosslinking at 65C. 10ul 0.5M EDTA, 20ul 1M Tris-HCl pH6.5 and 1ul proteinase K (20mg/ml) were added to de-crosslinked product and incubated for 1 hour at 45C. DNA was isolated by using Qiaquick PCR purification kit (Qiagen 28104). Libraries were prepared using the NEBNext® ChIP-seq Library Prep Master Mix Set for Illumina ® (NEB, E6240L) and QC'd using Agilent Technologies 2200 TapeStation to determine fragment size and PicoGreen (Life Technologies/Invitrogen, P7589) to quantify the concentration. Samples were pooled and submitted for SE50 sequencing using a HiSeq 2500.

**Epigenomic data analysis**

Reads were trimmed for both quality and Illumina adapter sequences using 'trim_galore' then aligned to mouse assembly mm9 with bowtie2 using the default parameters. Aligned reads with the same start site and orientation were removed using the Picard tool MarkDuplicates (http://broadinstitute.github.io/picard). ChIP density profiles were created by extending each read to the average library fragment size and then computing density using the BEDTools suite (http://bedtools.readthedocs.io). Reads were not extended when generating ATAC-seq read density. Enriched regions were discovered using MACS2 and scored against matched input libraries (fold change > 2 and p-value < 0.005). Peaks were then filtered against genomic 'blacklisted' regions (http://mitra.stanford.edu/kundaje/akundaje/release/blacklists/mm9-mouse/mm9-blacklist.bed.gz) and filtered peaks within 500 bp were merged to create a full peak atlas. Raw read counts were tabulated over this peak atlas using featureCounts (http://subread.sourceforge.net). All genome browser tracks and read density tables were normalized to a sequencing depth of ten million mapped reads. Peaks were annotated using linear genomic distance, with a gene assigned to a peak if it was within 50 kb upstream or downstream of the gene start or end, respectively. Motif signatures were obtained using the 'de novo' approach in Homer v4.5 (http://homer.ucsd.edu). Rescued peaks were defined as Stag2 regions in which Stag1 was enriched above input in the Stag2 KO, whereas non-rescued peaks showed no evidence of Stag1 binding in the Stag2 KO background.

**Hi-C**

Hi-C was performed as previously described(*66*). Briefly, $Lin^-$ bone marrow cells ($5x10^6$) were cross-linked in 1% formaldehyde for 10 minutes and quenched in 125mM glycine. Cross-linked cells were lysis in (10mM Tris-HCl pH8.0, 10mM NaCl, 0.2% Igepal CA-630 and Halt protease inhibitors Thermo Fisher 78429). After disruption, chromatin was solubilized and digested using 400Units of DpnII at 37°C overnight. DNA overhangs were then filled in with biotin-14-dATPs and ligated with T4 ligases at 16°C for 4 hours. Cross-links of ligated DNAs were reversed with proteinase K (Life Technologies, 25530-031) at 65°C overnight and purified using phenol:cholorform. After removal of biotin from unligated end, DNA was fragmented to 150-350 bps using an E220 sonicator (Covaris). After end repair, biotinylated DNA was collected using streptavidin beads (MyOne C1 beads, Life Technologies, 650.01) to prepare Hi-C libraries using the Illumina TruSeq Nano DNA kit. Hi-C libraries were sequenced on an Illumina HiSeq-4000 and raw sequencing data in the Fastq format were obtained.

Hi-C data was processed based on the previous method(*67*). Fastq files were mapped binned using c-World pipeline from the Dekker lab, which is available at a GitHub repository(*68*), https://github.com/dekkerlab/cMapping; https://github.com/dekkerlab/balance; https://github.com/dekkerlab/cworld-dekker). Briefly, 50bp paired end reads were truncated to 25bp starting at the 5-prime and then were iteratively mapped onto mm9. Uniquely mapped, paired end reads were collected and assigned to Dpn II restriction fragments based on their 5-prime locations. Mapped reads with same fragment ends and uniqueness were kept, and PCR duplicates were removed. Interaction heat-maps, insulation scores and loop pile-up were generated using scripts included in c-world pipeline. The loop coordinates used for loop pileup analysis were obtained from the mouse CH12.LX line(*12*). To increase resolution of Hi-C interaction profiles, intermediate validpairs files from the same genotype mice were pooled together using cooltools from the Mirny Lab (https://github.com/mirnylab). The insulation score change (ISC) between *Stag2*$^{-/y}$ and *Stag2*$^{fl/y}$ was calculated using 10kb binned Hi-C data and the ISC bins containing Stag2 rescued or non-rescued peaks were identified with Bedtools and annotated using ChIPseeker(*69*).

**Supplementary Figures:**

### Supplemental Fig. 1

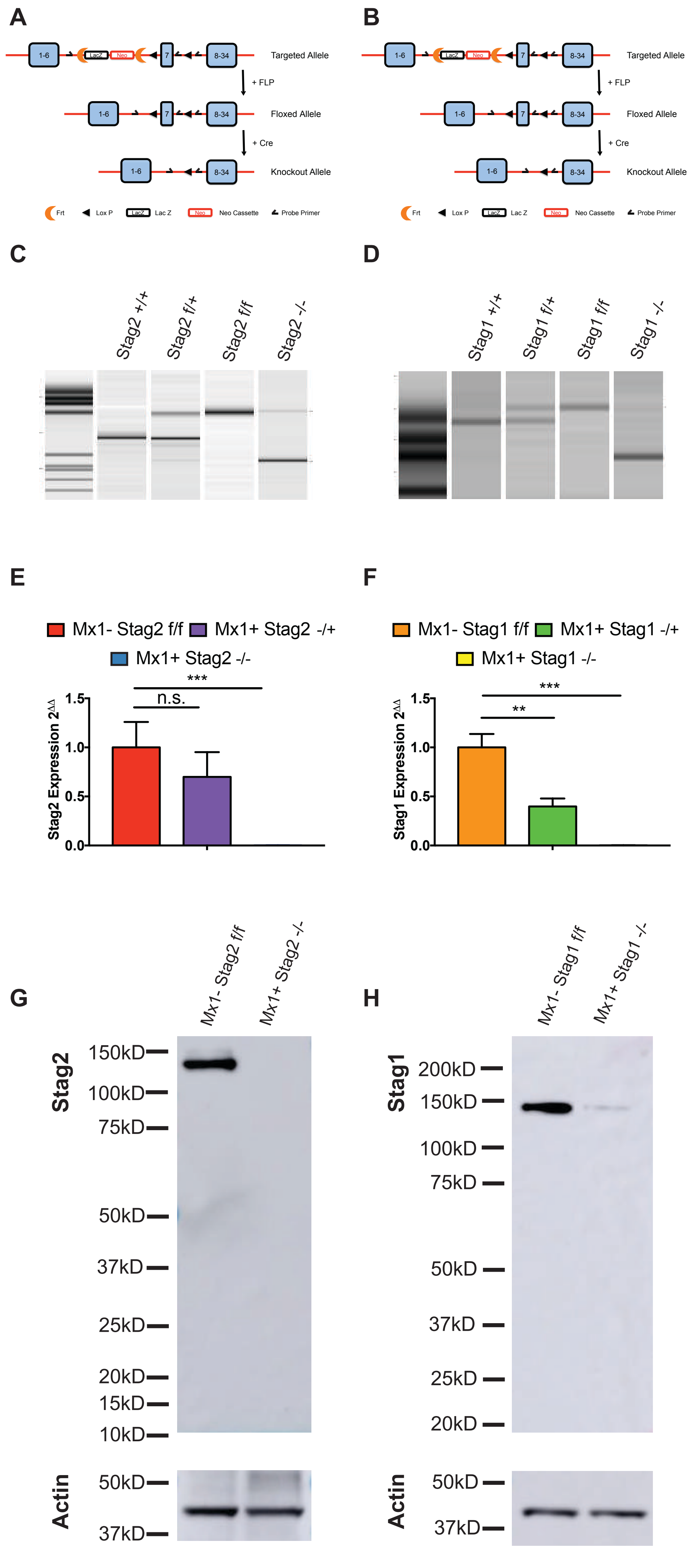

**Supplemental Fig 1.** Schematic representing the conditional knockout allele for **A)** Stag2 and **B)** Stag1 with each containing LoxP sites flanking exon 7. Digital PCR of blood genotyped for **C)** Stag2 or **D)** Stag1 from WT, floxed, and excised genotypes. **E)** RT-PCR measuring gene expression of Stag2 or **F)** Stag1 in floxed, heterozygous, and homozygous deletion. **G)** Full length western blot for Stag2 and **H)** Stag1 in floxed and excised bone marrow.

### Supplemental Fig. 2

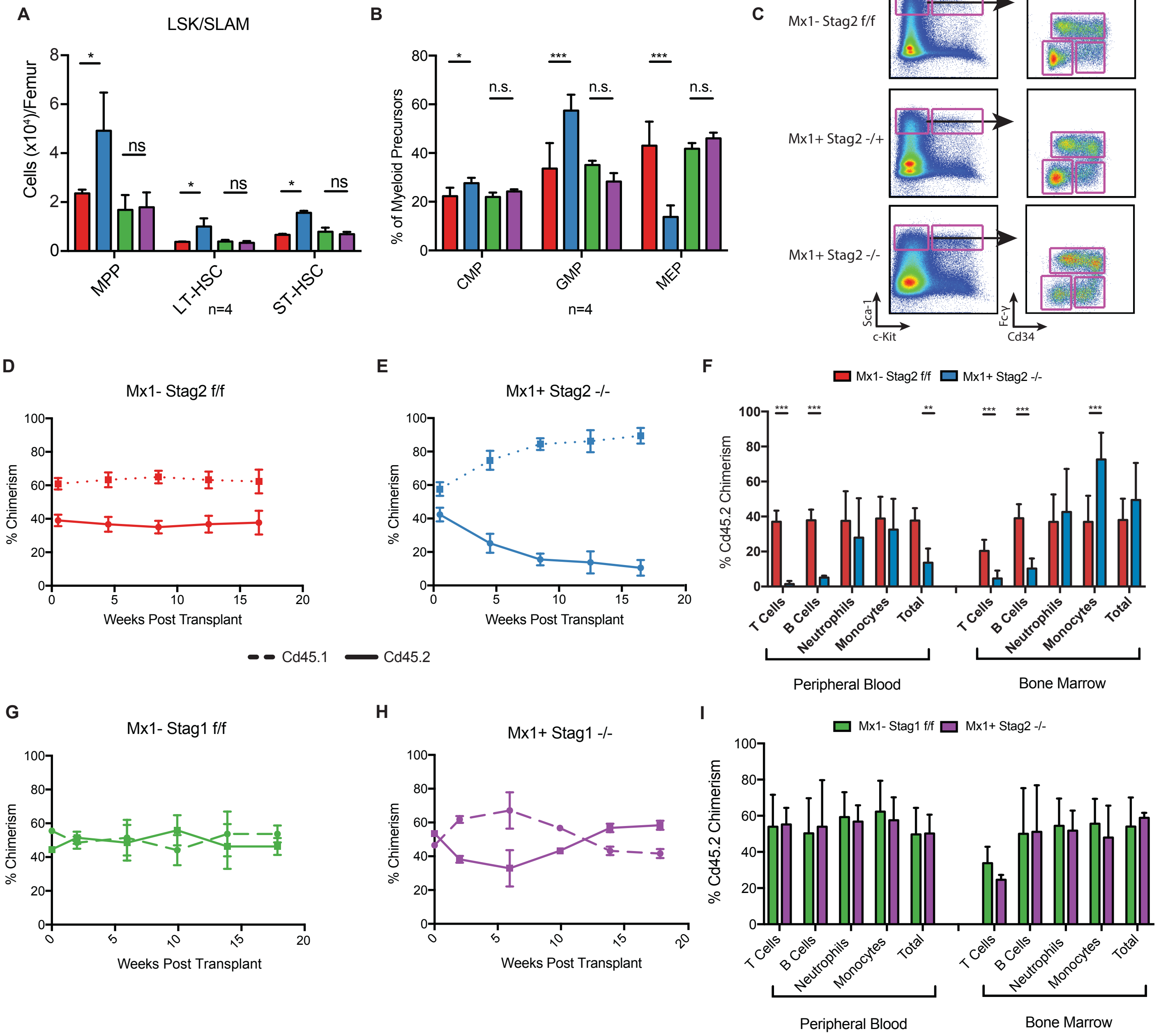

**Supplemental Fig 2. A)** Stag2 KO, but not Stag1 KO mice have increased hematopoietic stem cells as enumerated for each of the SLAM populations MPP, LT-HSC, and ST-HSC. **B)** Myeloid Progenitors ($Lin^{-}Kit^{+}Sca1^{-}$) show expanded granulocyte-macrophage precursors (GMP) at the expense of megakaryocyte-erythroid precursors (MEP). **C)** Representative flow cytometry scatter-plots of Stag2 floxed, heterozygous, and KO bone marrow (Parent gate is $Lin^{-}$ live singlets) showing increased LSK. Myeloid progenitors are gated by Cd34 and Fc-$\gamma$ revealing increased granulocyte-macrophage precursors and reduced megakaryocyte-erythroid progenitors. Competitive bone marrow transplantation of **D)** Stag2 WT or **E)** KO bone marrow mixed 1:1 with Cd45.1 normal marrow. Mice were injected with PIPC following engraftment at week 2. Flow cytometry of peripheral blood measured Cd45.2 chimerism every 4 weeks and **F)** at 16 weeks in the bone marrow. Competitive bone marrow transplantation of **G)** Stag1 WT or **H)** KO bone marrow mixed 1:1 with Cd45.1 normal marrow shows no difference in chimerism in peripheral blood **I)** or in the bone marrow at 16 weeks.

### Supplemental Figure 3

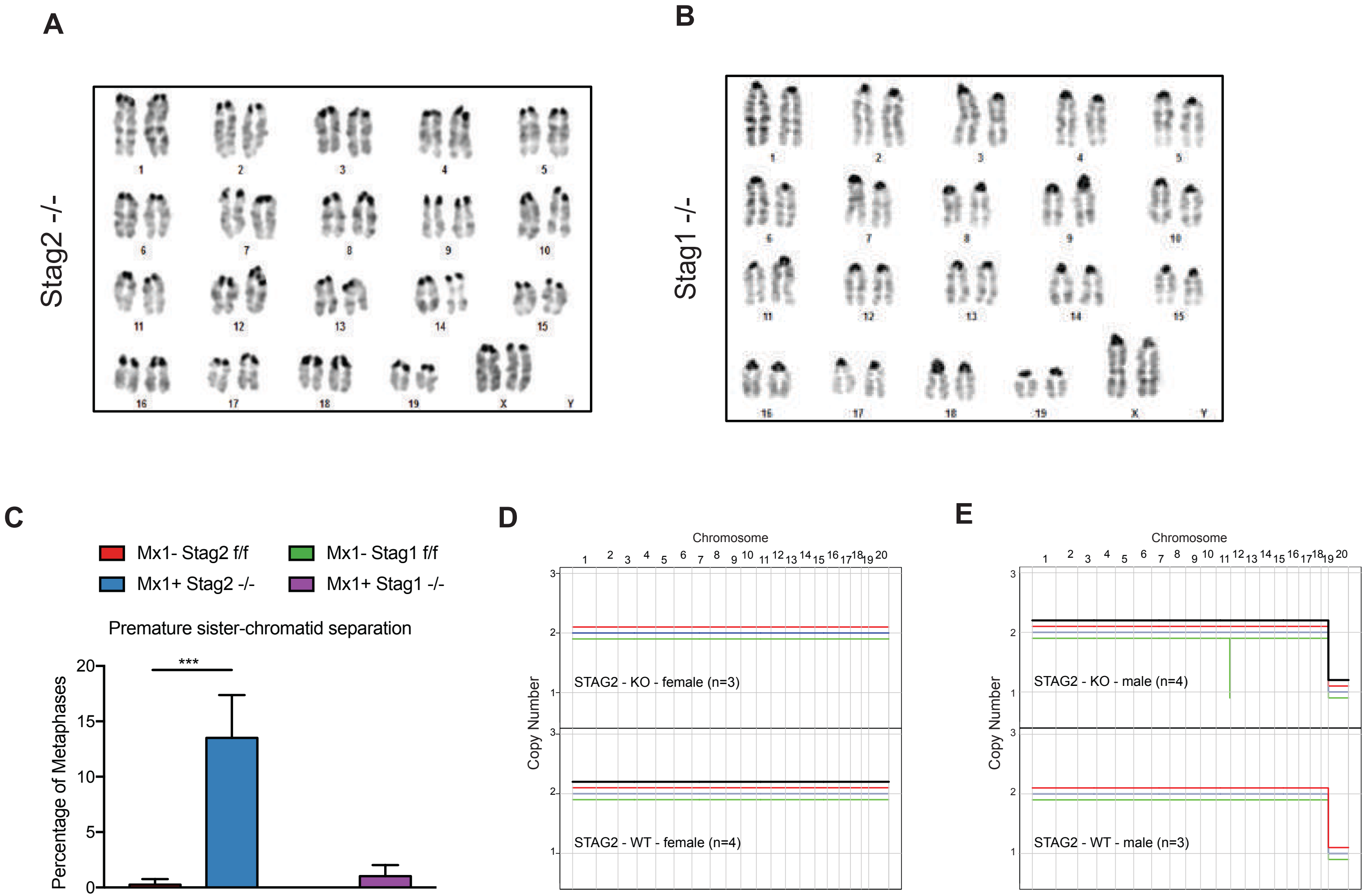

**Supplemental Fig 3. A)** Cytogenetic analysis of Stag2 and **B)** Stag1 KO mice. Representative metaphase spreads depicted for each genotype. All metaphase karyotypes were 40,XX [20] for Stag2 WT (n=4), Stag2 KO (n=3), Stag1 KO (n=3), and 40,XY [20] for Stag2 WT (n=3) and Stag2 KO (n=4). One of three Stag1 WT samples had a single tetraploid metaphase (40,XX[19], 80,XXXX [1]), and normal 40,XX [20] in the remaining two samples. **C)** Morphologic analysis of 100 metaphase figures for the presence of premature sister chromatid separation shows that more Stag2 KO cells had premature sister chromatid separation (mean=13.5%, p<0.001). **D)** Low depth whole genome sequencing of Stag2 WT and KO bone marrow for copy number shows no alterations genome wide in Stag2 WT females (n=4), Stag2 KO females (n=3), or **E)** Stag2 WT males (n=3). One of four Stag2 KO males shows a small 2Mb deletion at chr11qE2, and 3 have no alterations genome wide.

### Supplemental Fig. 4

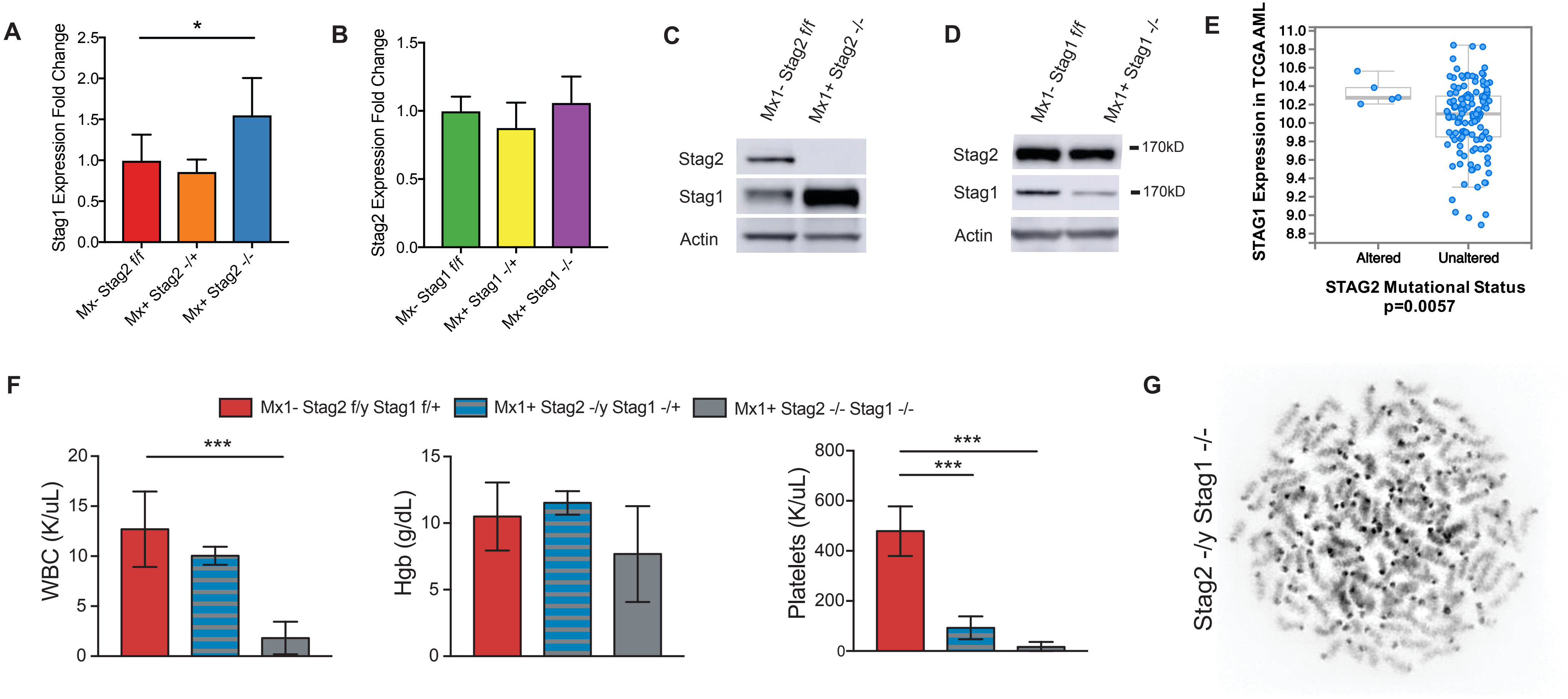

**Supplemental Fig 4. A-B)** RT-PCR measuring gene expression of (A) Stag1 or (B) Stag2 in floxed, heterozygous, and homozygous deletion of the opposing Stag gene. Stag1 expression increases in Stag2 KO bone marrow (p=0.1), but Stag2 expression does not change with Stag1 deletion. Western blot for Stag2 and Stag1 in **C)** Stag2 and **D)** Stag1 floxed and excised bone marrow. **E)** Patients with acute myeloid leukemia from The Cancer Genome Atlas (TCGA) with STAG2 mutations have higher levels of STAG1 expression compared to patients without cohesin mutations (p=0.006). **F)** Peripheral blood counts of Stag2 WT and KO mice with heterozygous or homozygous co-deletion of Stag1 taken at 7 days following PIPC (at the time Stag2/Stag1 KO mice were moribund) reveal decreased leukocytes in Stag2/Stag1 KO (p=0.001) and platelets in both Stag2 KO/Stag1$^{-/+}$ (p<0.001) and Stag2/Stag1 KO (p<0.001). **G)** Representative metaphase spreads depicted for male Stag2/Stag1 KO with chromosomal catastrophe

### Supplemental Fig. 5

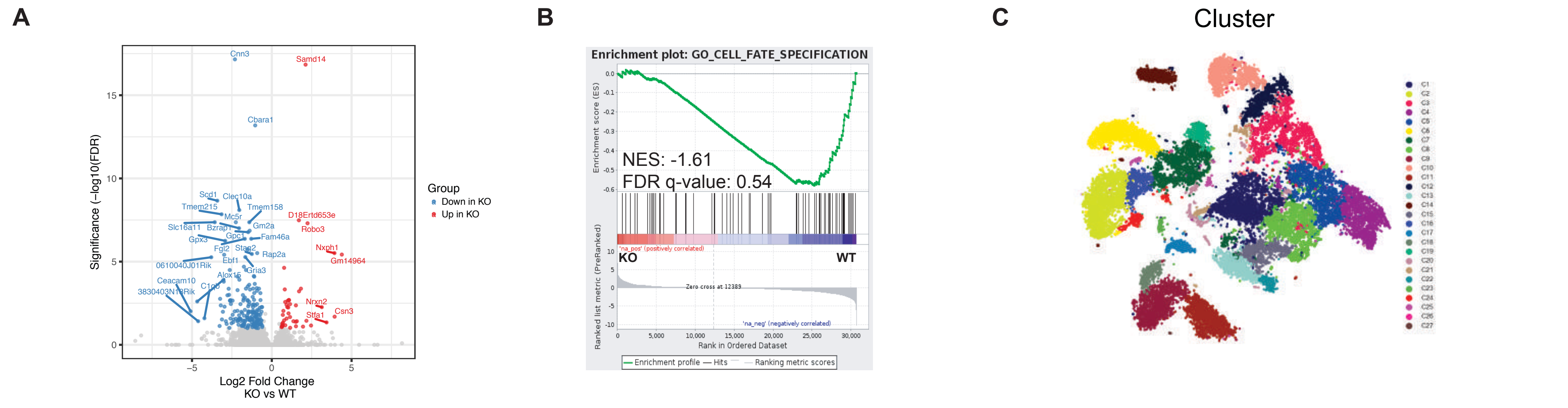


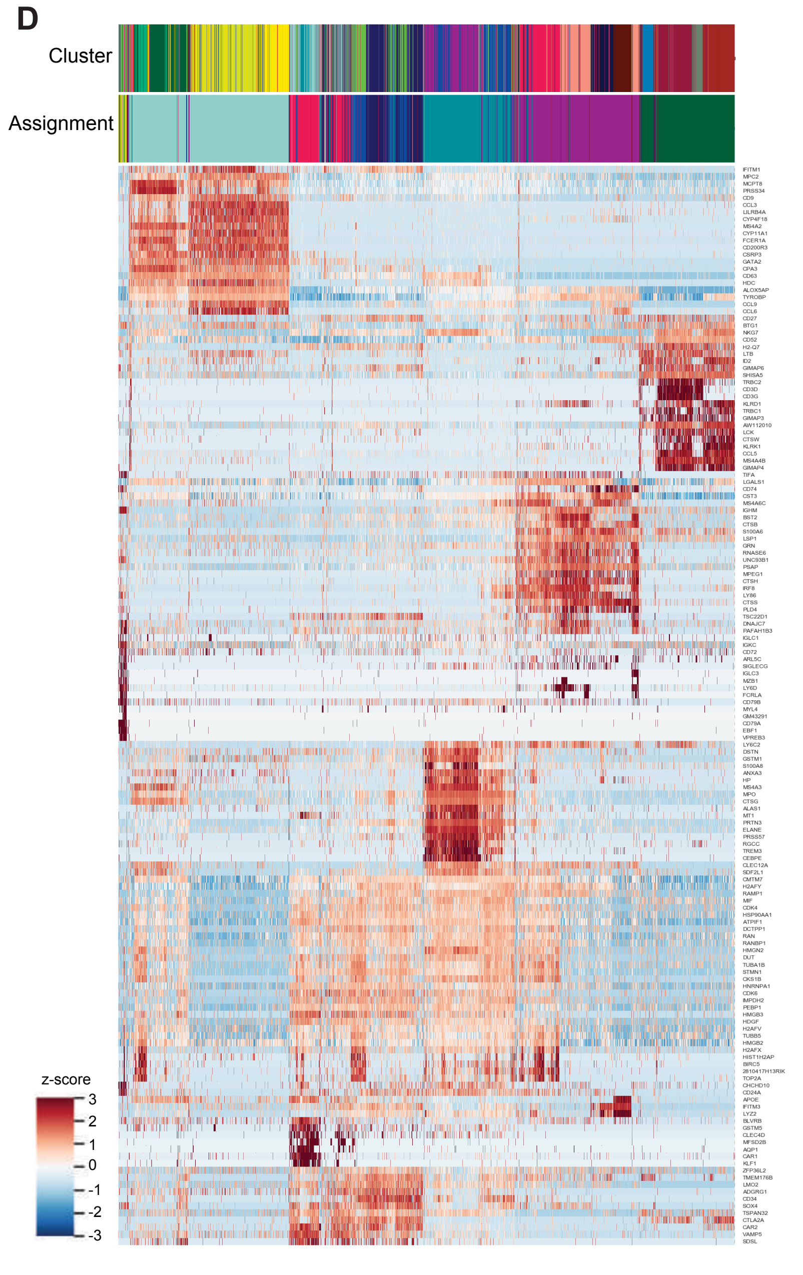


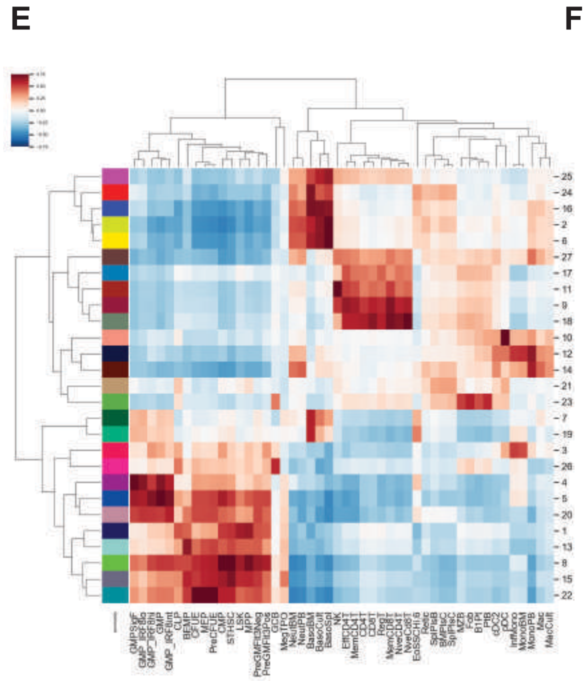


F

##### Positive Enrichment

| Gene Ontology | Name | NES | FDR q-value |
|---|---|---|---|
| GO_0034728 | NUCLEOSOME ORGANIZATION | 3.9097435 | 0 |
| GO_0006323 | DNA PACKAGING | 3.8820088 | 0 |
| GO_0065004 | PROTEIN-DNA COMPLEX ASSEMBLY | 3.5663702 | 0 |
| GO_0006334 | NUCLEOSOME ASSEMBLY | 3.5006585 | 0 |
| GO_0031497 | CHROMATIN ASSEMBLY | 2.8557205 | 0 |
| GO_0006333 | CHROMATIN ASSEMBLY OR DISASSEMBLY | 2.6403115 | 0 |
| GO_0007059 | CHROMOSOME SEGREGATION | 2.521101 | 0 |

##### Negative Enrichment

| Gene Ontology | Name | NES | FDR q-value |
|---|---|---|---|
| GO_0042113 | B CELL ACTIVATION | 2.9033923 | 0 |
| GO_0030098 | LYMPHOCYTE DIFFERENTIATION | 2.5199957 | 0 |
| GO_0042110 | T CELL ACTIVATION | 2.3701916 | 0 |
| GO_0030217 | T CELL DIFFERENTIATION | 2.235279 | 0 |
| GO_0002521 | LEUKOCYTE DIFFERENTIATION | 2.2332351 | 0 |
| GO_0044087 | REGULATION OF CELLULAR COMPONENT BIOGENESIS | 2.0384235 | 0 |
| GO_0042113 | B CELL ACTIVATION | 2.9033923 | 0 |

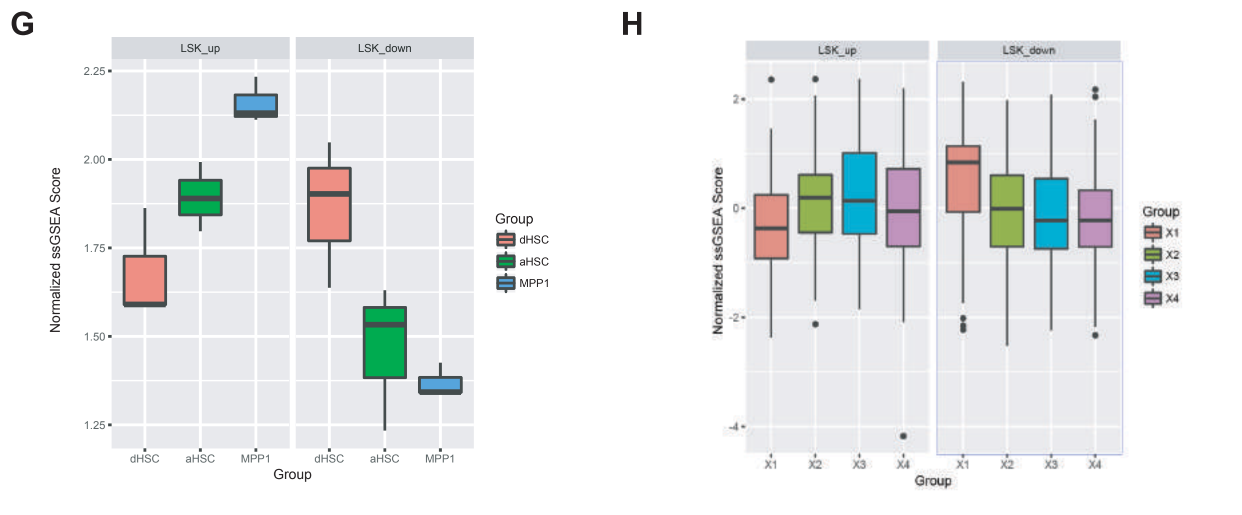

**Supplemental Fig 5. A)** Volcano plot for differentially expressed genes by RNA sequencing in LSK cells of Stag2 WT and Stag2 KO. Genes decreased in expression in Stag2 KO in blue (n=186) and genes increased in expression in Stag2 KO in red (n=42). **B)** Gene-set enrichment analysis of LSK RNAseq shows decreased expression of the GO Cell Fate Specification gene set. **C**) t-SNE projection of library-size normalized and log transformed data for complete collection (24,153 cells). Each dot represents a single cell colored by Phenograph clustering(*61*). **D**) Pearson correlation between centroids of Phenograph clusters to standardized bulk RNA-sequencing data from selected sorted mouse hematopoietic cells populations (from Haemopedia-Mouse RNAseq(*62*)). **E**) Heatmap of normalized and log transformed expression of top 20 differentially expressed genes per inferred lineage; cells and genes are hierarchically clustered. Differential expression was determined by calculating, for every gene, the Wasserstein distance between normalized and log-transformed expression in cells from inferred lineage and all other cells; top heatmaps show assignment of cells to clusters and inferred lineages labeled in Figure 2D. **F)** Gene set enrichment meeting threshold of NES >2, FDR <0.25 for positive enrichment (top) and negative enrichment (bottom) with gene ontology. **G)** Single cell RNA seq data from Cabezas- Wallscheid et al. Genes increased in Stag2 KO LSK are enriched for genes expressed in active HSC and downregulation of quiescent genes. **H)** Comparisons to single cell RNAseq further refines the decreased expression in stage 1 of the dHSC (X1) compared to stages 2-4 (vs X2 $p=2.8 \times 10^{5}$; vs X3 $p=1.9 \times 10^{5}$; vs X4 $p=2.5 \times 10^{6}$). Statistical comparisons were generated using pairwise Student's t-test.

### Supplemental Fig. 6

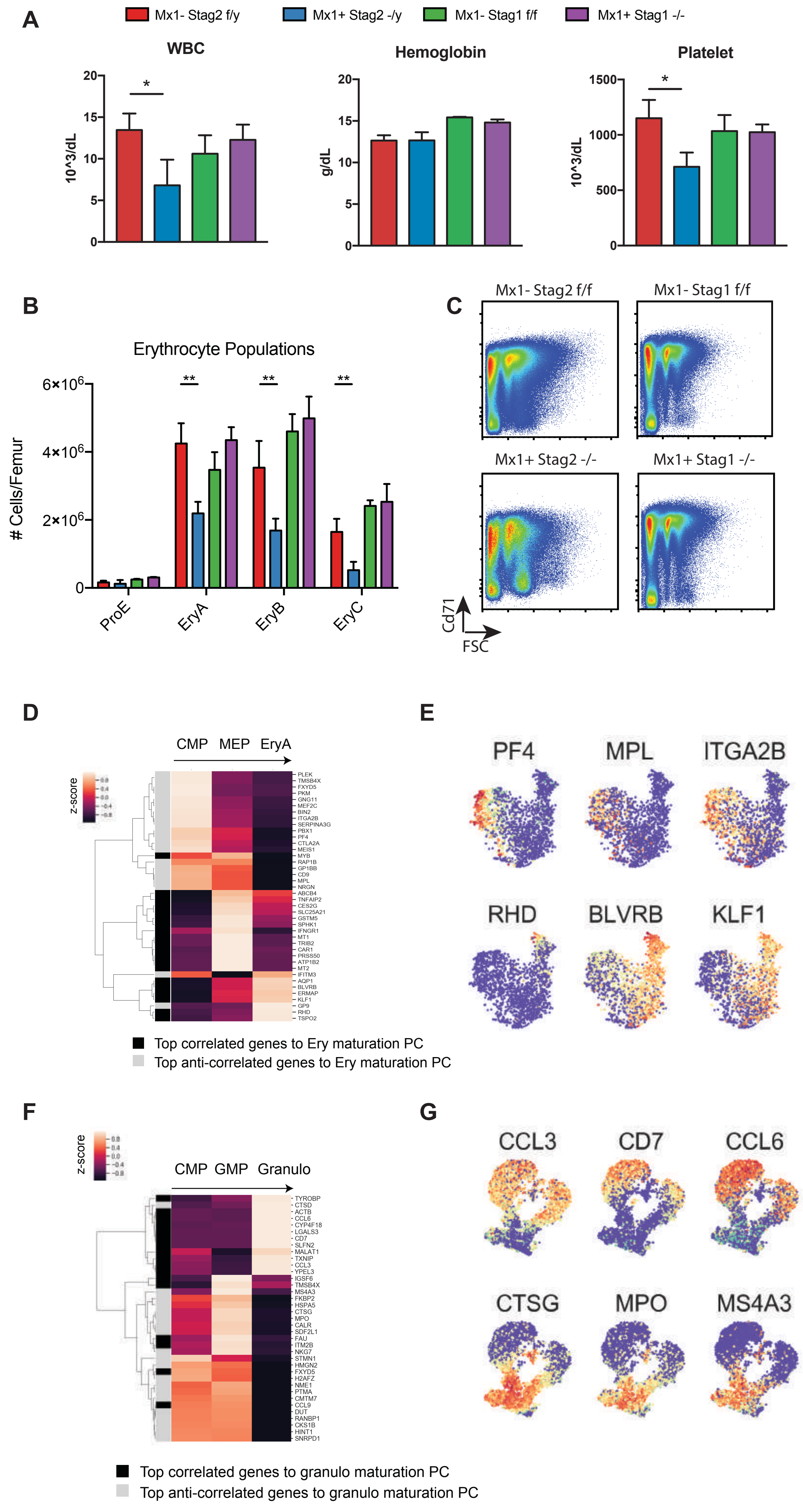

**Supplemental Fig 6. A)** Peripheral blood counts of Stag2 and Stag1 WT and KO mice taken at 8 weeks following PIPC treatment reveal decreased leukocytes (p=0.02) and platelets (p=0.01) in Stag2 KO mice and no changes in Stag1 KO mice. No differences in hemoglobin were seen in either Stag2 or Stag1 KO. **B)** Flow cytometric analysis of erythroid development using Ter-119 and Cd71 showing reduced mature erythroid populations (EryA p=0.002; EryB p=0.007; EryC p=0.005) in Stag2 KO bone marrow. **C)** Representative flow cytometry scatter-plots of Stag2 WT / KO and Stag1 WT / KO bone marrow showing Stag2 KO cells fail to decrease in FSC during maturation (Parent Gate on Ter-119$^{+}$ Cells). **D**) Heatmap of bulk RNA-sequencing data for cells with progressive erythroid lineage commitment (from GSE60101(*70*)) showing normalized and standardized expression of genes most correlated or anticorrelated with erythroid maturation component. Correlations were computed using the Pearson method. **E**) t-SNE projection of library-size normalized and log transformed data for inferred MEP subset (1787 cells). Each dot represents a single cell colored by expression of labelled genes. **F**) Heatmap of bulk RNA-sequencing data for cells with progressive granulocyte lineage commitment (from GSE60101(*70*)) showing normalized and standardized expression of genes most correlated or anticorrelated with granulocyte maturation component. Correlations were computed using the Pearson method. **G**) t-SNE projection of library-size normalized and log transformed data for inferred granulocyte subset (6316 cells). Each dot represents a single cell colored by expression of labelled genes.

### Supplemental Fig. 7

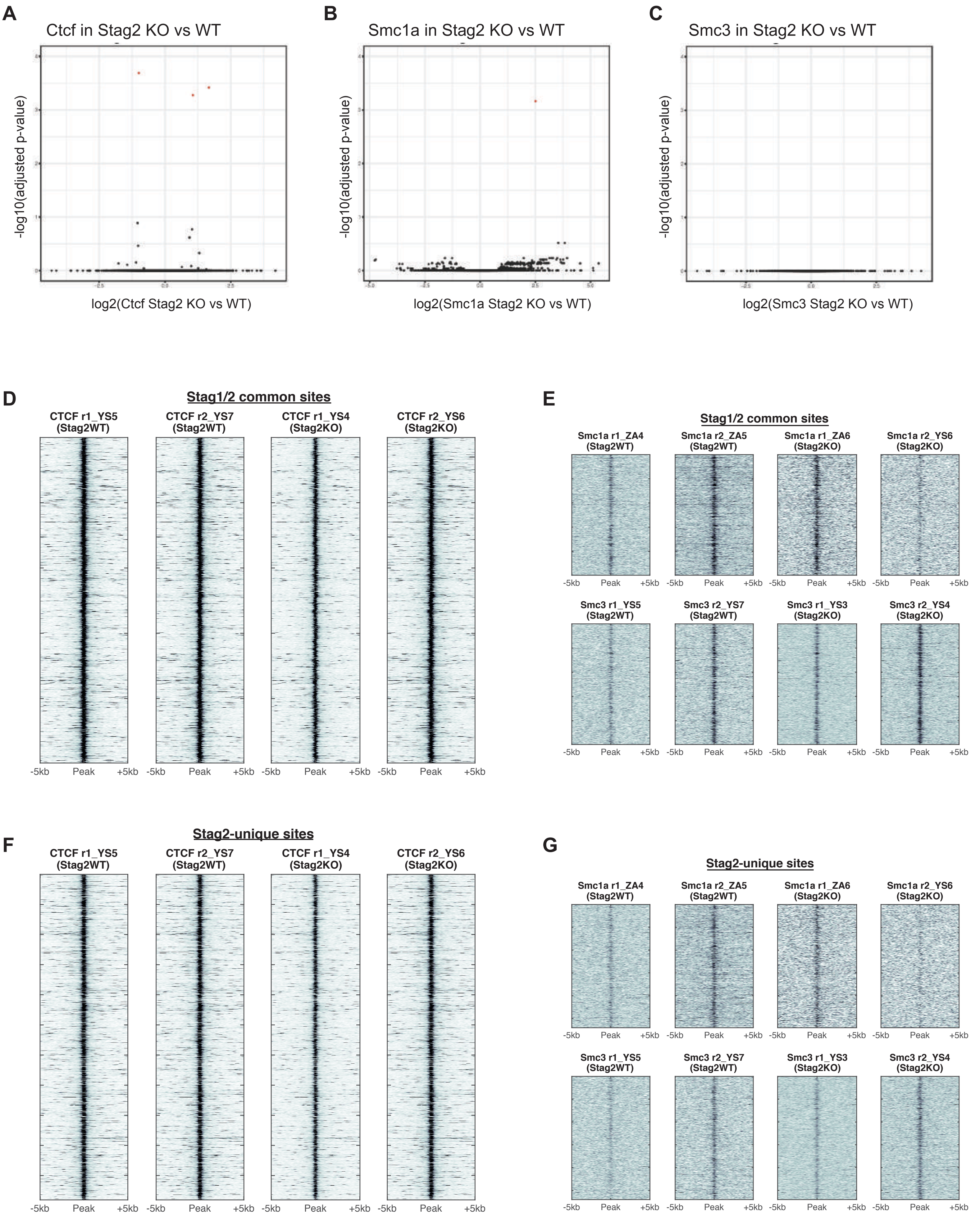

**Supplemental Fig 7.** Chromatin immunoprecipitation and sequencing for **A)** Ctcf **B)** Smc1a and **C)** Smc3 in Stag2 WT (n=2) and KO (n=2) HSPC. Volcano plots show lack of statistically significant differential loci. Heatmaps for **D)** Ctcf and **E)** Smc1a (top row)/Smc3 (bottom row) at Stag2/Stag1 commonly bound sites. Heatmaps for **F)** Ctcf and **G)** Smc1a (top row)/Smc3 (bottom row) at Stag2-uniquely bound sites. No differential occupancy in either Stag1/2 common or Stag2-unique sites were identified.

### Supplemental Fig. 8

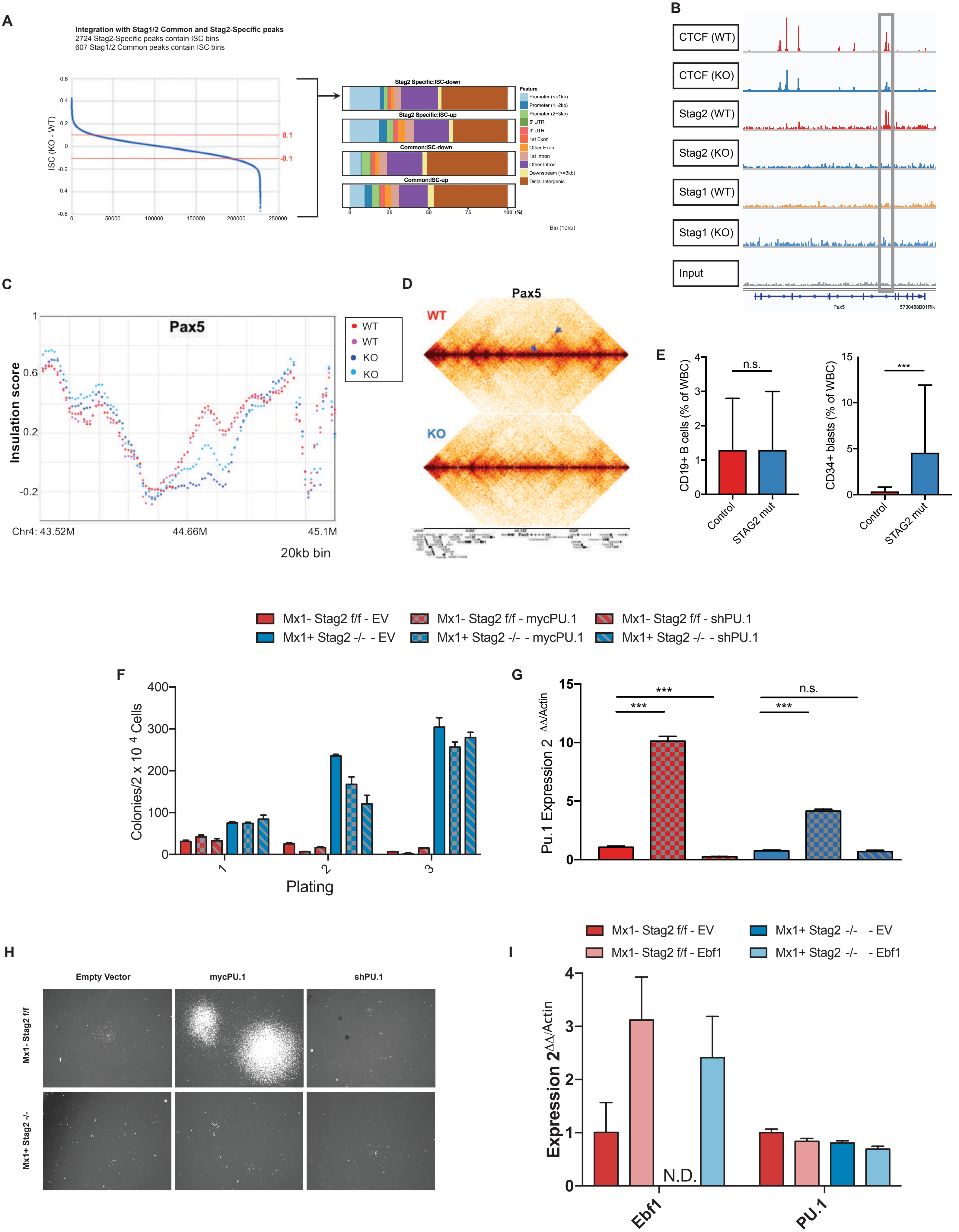

**Supplemental Fig 8. A)** Insulation score changes (ISC: Insulation score KO – Insulation score WT) from Hi-C analysis of Stag2 WT and KO lineage negative bone marrow using windows of 20kB. Stacked bar plot of genomic feature for common and Stag2-specific sites according to gain or loss of insulation. **B)** IGV track of the Pax5 locus with Stag2 and Ctcf binding at a single locus that is lost in Stag2 KO and not bound by Stag1 either in WT or KO. **C)** ISC plotted across Pax5 for Stag2 WT (n=2; shades of red) and KO (n=2; shades of blue) shows marked loss of insulation. **D)** Contact map of Pax5 shows loss of cis-interaction at two loci (arrows).
**E)** Enumeration of mature B cells (CD34-CD19+) and CD34+ blasts in STAG2 mutated MDS patients (n=11) compared to controls (n=15). **F)** Methylcellulose colony assay using IL-3, SCF, and IL-6 enriched media for stem cell replating. Stag2 WT and KO marrow were infected with lentivirus containing GFP-tagged empty vector, GFP-mycPu.1, or GFP-shPu.1. 20,000 $GFP^+$ cells were serially replated with overexpression and shRNA manipulation of Pu.1 unable to abrogate serial replating of the Stag2 KO cells. **G)** RT-PCR for Spi1 shows lower expression in Stag2 KO compared to Stag2 WT (p=0.01). Spi1 expression increased 10-fold in Stag2 WT mycPu.1 transfected cells (p<0.001), and a 4-fold reduction was observed in Stag2 WT shPu.1 transfected cells (p<0.001). Spi1 expression increased 5-fold in Stag2 KO mycPU.1 transfected cells (p<0.001). The further decrease of Spi1 expression in Stag2 KO shPu.1 was not statistically significant. **H)** B-cell colony formation was markedly reduced in EV-Stag2 KO compared to EV-Stag2 WT (p=0.01). Compared to EV, colony number was reduced with shRNA knockdown of Pu.1 in both Stag2 WT (p=0.09) and Stag2 KO (p=0.09) but did not reach statistical significance. B-cell colonies were markedly larger and of higher cell output in Stag2 WT mycPu.1 transfected cells than in any other group. **I)** RT-PCR for Ebf1 shows higher expression in both Stag2 WT and Stag2 KO cells transfected with Ebf1. Spi1 expression did not change.
